# The structural, dynamical and biochemical characterizations of *Verticillium dahliae* pectate lyase, VdPelB, highlight its specificities

**DOI:** 10.1101/2022.11.09.515409

**Authors:** Josip Safran, Vanessa Ung, Julie Bouckaert, Olivier Habrylo, Roland Molinié, Jean-Xavier Fontaine, Adrien Lemaire, Aline Voxeur, Serge Pilard, Corinne Pau-Roblot, Davide Mercadante, Jérôme Pelloux, Fabien Sénéchal

**Affiliations:** UMRT INRAE 1158 BioEcoAgro – BIOPI Biologie des Plantes et Innovation, Université de Picardie, 33 Rue St Leu, 80039 Amiens, France; School of Chemical Sciences, The University 9 of Auckland, Private Bag 92019, Auckland 1142, New Zealand; UMR 8576 Unité de Glycobiologie Structurale et Fonctionnelle (UGSF) IRI50, Avenue de Halley, 59658 Villeneuve d’Ascq, France; Université Paris-Saclay, INRAE, AgroParisTech, Institut Jean-Pierre Bourgin (IJPB), 78000, Versailles, France; Plateforme Analytique, Université de Picardie, 33, Rue St Leu, 80039 Amiens, France

**Keywords:** Pectate lyase, pectins, homogalacturonan, oligogalacturonides, Flax, *Verticillium dahliae*.

## Abstract

Pectins, complex polysaccharides and major components of the plant primary cell wall, can be degraded by pectate lyases (PLs). PLs cleave glycosidic bonds of homogalacturonans (HG), the main pectic domain, by β-elimination, releasing unsaturated oligogalacturonides (OGs). To understand the catalytic mechanism and structure/function of these enzymes, we characterized VdPelB from *Verticillium dahliae*, a plant pathogen. We first solved the crystal structure of VdPelB at 1.2Å resolution showing that it is a right-handed parallel β-helix structure. Molecular dynamics (MD) simulations further highlighted the dynamics of the enzyme in complex with substrates that vary in their degree of methylesterification, identifying amino acids involved in substrate binding and cleavage of non-methylesterified pectins. We then biochemically characterized wild type and mutated forms of VdPelB. VdPelB was most active on non-methylesterified pectins, at pH 8 in presence of Ca^2+^ ions. VdPelB-G125R mutant was most active at pH 9 and showed higher relative activity compared to native enzyme. The OGs released by VdPelB differed to that of previously characterized PLs, showing its peculiar specificity in relation to its structure. OGs released from *Verticillium-*partially tolerant and sensitive flax cultivars differed which could facilitate the identification VdPelB-mediated elicitors of defence responses.

## 1. Introduction

Primary cell wall, a complex structure of proteins and polysaccharides, cellulose and hemicelluloses, is embedded in a pectin matrix. Pectins, are complex polysaccharides composing up to 30% of cell wall dry mass in dicotyledonous species [1]. Pectin is mainly constituted of homogalacturonan (HG), rhamnogalacturonan I (RG-I) and rhamnogalacturonan II (RG-II) domains, but its composition can differ between plant organs and among species. The most abundant pectic domain is HG, a linear homopolymer of α-1,4-linked galacturonic acids (GalA), which represents up to 65% of pectins [2]. During synthesis, HG can be O-acetylated at O-2 or O-3 and/or methylesterified at C-6 carboxyl, before being exported at the cell wall with a degree of methylesterification (DM) of ∼80% and a degree of acetylation (DA) of ∼5-10%, depending on species [3]. At the wall, HG chains can be modified by different enzyme families, including pectin acetylesterase (PAEs; EC 3.1.1.6), pectin methylesterases (PMEs; CE8, EC 3.1.1.11), polygalacturonases (PGs; GH28, EC 3.2.1.15, EC 3.2.1.67, EC 3.2.1.82), and pectin lyases-like (PLLs), which comprise pectate lyases (PLs; EC 4.2.2.2) and pectin lyase (PNLs, EC 4.2.2.10). All these enzymes are produced by plants to fine-tune pectin during development [4–8], but they are also secreted by most phytopathogenic bacteria and fungi during plant infection [9–13]. PMEs and PAE hydrolyse the O6-ester and O2-acetylated linkages, respectively, leading to a higher susceptibility of HG to PG- and PLL-mediated degradation [14]. PLL are pectolytic enzymes that cleave HG via a β-elimination mechanism leading to the formation of an unsaturated C4-C5 bond [15], and can be divided into two subfamilies depending on their biochemical specificities: i) PLs have a high affinity for non- or low-methylesterified pectins and an optimal pH near 8.5. Their activity requires Ca^2+^ ions. ii) PNLs are most active on high-DM pectins at acidic pH values [16]. Both type of enzymes can degrade HG chains, and release oligogalacturonides (OGs), but their mode of action can differ. For PLs both endo and exo modes of action have been described, while only endo-PNL have been characterised so far [17]. For the latter, it was notably shown that endo-PNLs from *B. fuckeliana*, *A. parasiticus* and *Aspergillus* sp., can first release OGs with degrees of polymerisation (DP) 5–7, that are subsequently used as substrates, generating OGs of DP3 and DP4 as end-products [9,18,19]. Despite having the same DP, the final products can differ in their degrees and patterns of methylesterification and acetylation (DM/DA) depending on the enzymes’ specificities; implying potential differences in substrate binding, and therefore in PLLs fine structures. Several crystallographic structures of bacterial and fungal PLL have been reported [15,20–24]. Overall, the PLL fold resemble that of published PME, PGs and rhamnogalacturonan lyases [25–27], and is composed of three parallel β-sheets forming a right-handed parallel β-helix. The three β-sheets are called PB1, PB2 and PB3 and the turns connecting them T1, T2 and T3 [28]. The active site features three Asp, localized on the PB1 β-sheet and, in the case of PLs, accommodates Ca^2+^ [29]. In PNLs, Ca^2+^ is, on the other hand, replaced by Asp [15]. Additionally, the PL binding site is dominated by charged acidic and basic residues (Gln, Lys, Arg) which can accommodate negatively charged pectate substrates. In contrast, the PNL binding site is dominated by aromatic residues [15, 29], which have less affinity for lowly methylesterified pectins. These differences in structure could translate into distinct enzyme dynamics when in complex with substrates of varying degrees of methylesterification.

In fungi, PLLs are encoded by large multigenic families which are expressed during infection. *Verticillium dahliae* Kleb., a soil-borne vascular fungus, targets a large number of plant species, causing Verticillium wilt disease to become widespread among fiber flax, with detrimental effects to fiber quality [30–32]. *V. dahliae* infects plants by piercing the root surface using hyphae, secreting a number of pectinolytic enzymes, including thirteen PLLs. Considering the role of PLLs in determining pathogenicity, it is of paramount importance to determine their biochemical and structural properties [30,33,34]. This could allow engineering novel strategies to control or inhibit, the pathogen’s pectinolytic arsenal. For this purpose, we characterized, via combined experimental and computational approaches, one *V*. *dahliae* PLL (VdPelB, VDAG_04718) after its heterologous expression in *P. pastoris*. The obtention of the 3D structure of VdPelB after X-ray diffraction and the analysis of enzyme dynamics when in complex with substrates of distinct DM, allowed the identification of the residues favouring pectate lyase (PL) activity. Experiments confirmed the importance of these residues in mediating PL activity showing that VdPelB is a *bona fide* PL, that releases peculiar OG as compared to previously characterized PLs. More importantly, the OGs released from roots of *Verticillium-*partially tolerant and sensitive flax cultivars differed, paving the way for the identification of VdPelB-mediated OGs that can trigger plant defence mechanisms.

## 2. Material and methods

### 2.1. Bioinformatical analysis

*Verticillium dahliae* PLL sequences were retrieved using available genome database (ftp.broadinstitute.org/). SignalP-5.0 Server (http://www.cbs.dtu.dk/services/ SignalP/) was used for identifying putative signal peptide and putative glycosylation sites were predicted using NetNGlyc 1.0 Server (http://www.cbs.dtu.dk/services/NetNGlyc/) and NetOGlyc 4.0 Server (http://www.cbs.dtu.dk/services/NetOGlyc/). Sequences were aligned and phylogenetic analysis was carried out using MEGA multiple sequence alignment program (https://www.megasoftware.net/). Homology models were created using I-TASSER structure prediction software (https://zhanglab.ccmb.med.umich.edu/I-TASSER/) and UCSF Chimera (http://www.cgl.ucsf.edu/chimera/) was used for creation of graphics.

### 2.2. Fungal strain and growth

*V.dahliae* was isolated from CALIRA company flax test fields (Martainneville, France) and was kindly provided by Linéa-Semences company (Grandvilliers, France). Fungus was grown as described in Safran et al. [35]. Briefly, fungus was grown in polygalacturonic acid sodium salt (PGA, P3850, Sigma) at 10 g.L^−1^ and in pectin methylesterified potassium salt from citrus fruit (55–70% DM, P9436, Sigma) solutions to induce PLL expression. After 15 days of growth in dark conditions at 25°C and 80 rpm shaking, mycelium was collected and filtered under vacuum using Buchner flask. Collected mycelium was frozen in liquid nitrogen, lyophilized and ground. Isolation of RNA and cDNA synthesis was realized as previously described in Lemaire et al.[36].

### 2.3. Cloning, heterologous expression and purification of VdPelB

*V. dahliae* PelB coding sequence (VdPelB, UNIPROT: G2X3Y1, GenBank: EGY23280.1), minus the signal peptide was amplified using cDNA and gene-specific primers. VdPelB mutants were generated using cDNA and specific primers carrying mutations (Table S1). Cloning, heterologous expression in *P. pastoris* and purification of VdPelB was done as previously described in Safran et al. [35].

### 2.4. Crystallization of VdPelB

VdPelB was concentrated at 10 mg.mL^-1^ in Tris-HCl pH 7.5 buffer. Crystallization conditions were screened using the sitting-drop vapor-diffusion method. VdPelB (100 nL) was mixed with an equal volume of precipitant (1:1) using Mosquito robot (STP Labtech). The crystals that resulted in best diffraction data were obtained with 0.1 M MIB (Malonic acid, Imidazole, Boric acid system) at pH 8.0, with 25 % PEG 1500 as the precipitant (condition B5 from the PACT premier kit, Molecular Dimensions, Sheffield, UK) after 1 month. Optimization was realized using the hanging drop vapor-diffusion method forming the drop by mixing 1 µL of precipitant solution with 1 µL of the enzyme. The large beam-like crystals were cryo-protected by increasing PEG 1500 concentration to 35%, before mounting them in a loop and flash-cooling them in liquid nitrogen.

### 2.5. VdPelB X-ray data collection and processing

X-ray diffraction data were collected at the PROXIMA-2a beamline of the Soleil synchrotron (Saint Aubin, France), at a temperature of −173°C using an EIGER 9M detector (Dectris). Upon a first data collection to 1.3 Å resolution, three more data sets were collected from the same crystal in order to obtain a complete data set. Thereby the kappa angle was tilted once to 30°, once to 60° and finally a helical data set was collected at 1.2 Å resolution. The reflections of each data set were indexed and integrated using XDS [37], scaled and merged using XSCALE [38]. The VdPelB crystal has a primitive monoclinic lattice in the P 1 2_1_ 1 space group, with two molecules contained in asymmetric unit [39].

### 2.6. Structure solution and refinement

The structure of VdPelB was solved by molecular replacement using *Phaser* [40]. The data were phased using pectate lyase BsPelA (PDB: 3VMV, Uniprot D0VP31), as a search model [41]. Model was build using *Autobuild* and refined using *Refine* from PHENIX suite [42]. The model was iteratively improved with *Coot* [43] and *Refine.* The final structure for VdPelB has been deposited in the Protein Data Bank (PDB) as entry 7BBV.

### 2.7. VdPelB biochemical characterization

Pierce BCA Protein Assay Kit (Thermo Fisher Scientific, Waltham, Massachusetts, United States) was used to determine the protein concentration, with Bovine Serum Albumin (A7906, Sigma) as a standard. Deglycosylation was performed using Peptide-N-Glycosidase F (PNGase F) at 37 °C for one hour according to the supplier’s protocol (New England Biolabs, Hitchin, UK). Enzyme purity and molecular weight were estimated using a 12% SDS-PAGE and mini-PROTEAN 3 system (BioRad, Hercules, California, United States). Gels were stained using PageBlue Protein Staining Solution (Thermo Fisher Scientific) according to the manufacturer’s protocol.

The substrate specificity of VdPelB was determined using PGA (81325, Sigma) and citrus pectin of various DM: 20–34% (P9311, Sigma), 55–70% (P9436, Sigma) and >85% (P9561, Sigma), with 0.5 µM CaCl_2_ or 5 µM EDTA (Sigma) final concentrations. Enzyme activity was measured by monitoring the increase in optical density at 235 nm due to formation of unsaturated uronide product using UV/VIS Spectrophotometer (PowerWave Xs2, BioTek, France) during 60 min. The optimum temperature was determined by incubating the enzymatic reaction between 25 and 45°C for 8 min using PGA as a substrate (0.4%, w/v). The optimum pH was determined between pH 5 and 10 using sodium phosphate (NaP, pH 5 to 7) and Tris-HCl buffer (pH 7 to 10) and 0.4% (w/v) PGA as a substrate. All experiments were realized in triplicate.

### 2.8. Digestion of commercial pectins and released OGs profiling

OGs released after digestions by recombinant VdPelB or commercially available Aspergillus PL (named AsPel) were identified as described in Voxeur et al., 2019 [9], using a novel in-house OGs library. Briefly, DM 20–34% (P9311, Sigma) and sugar beet pectin with DM 42% and degree of acetylation (DA) 31% (CP Kelco, Atlanta, United States) were prepared at 0.4 % (w/v) final concentration diluted in 50 mM Tris-HCl buffer (pH 8) and incubated with either VdPelB or AsPel (E-PCLYAN, Megazyme). For each substrate, enzyme concentrations were adjusted to have enzymes at iso-activities (Table S2). For each substrate two dilutions, were used for analysing OGs released in early, VdPelB-2 and AsPel-2, and late phase, VdPelB-1 and AsPel-1, of digestions. Digestions were performed overnight. Non-digested pectins were pelleted by centrifugation and the supernatant dried in a speed vacuum concentrator (Concentrator plus, Eppendorf, Hamburg, Germany). Separation of OGs was done as previously described using an ACQUITY UPLC Protein BEH SEC column (125Å, 1.7 μm, 4.6 mm x 300 mm) [44]. The intensities were defined as the area under the curve, for each OG. Peak areas were clustered by hierarchical clustering with complete linkage on the euclidian distance matrix and visualized in the heatmap-package using R version 3.6.0.

### 2.9. Digestion of commercial pectins and released OGs profiling

OGs released after digestions by recombinant VdPelB or commercially available Aspergillus PL (named AsPel) were identified as described in Voxeur et al., 2019 [9], using a novel in-house OGs library. Briefly, DM 20–34% (P9311, Sigma) and sugar beet pectin with DM 42% and degree of acetylation (DA) 31% (CP Kelco, Atlanta, United States) were prepared at 0.4 % (w/v) final concentration diluted in 50 mM Tris-HCl buffer (pH 8) and incubated with either VdPelB or AsPel (E-PCLYAN, Megazyme). For each substrate, enzyme concentrations were adjusted to have enzymes at iso-activities (**Table S2**). For each substrate two dilutions, were used for analysing OGs released in early, VdPelB-2 and AsPel-2 and late phase, VdPelB-1 and AsPel-1, of digestion. Digestions were performed overnight. Non-digested pectins were pelleted by centrifugation and the supernatant dried in a speed vacuum concentrator (Concentrator plus, Eppendorf, Hamburg, Germany). Separation of OGs was done as previously described using an ACQUITY UPLC Protein BEH SEC column (125Å, 1.7 μm, 4.6 mm x 300 mm) [44]. The intensities were defined as the area under the curve, for each OG. Peak areas were clustered by hierarchical clustering with complete linkage on the euclidian distance matrix and visualized in the heatmap-package using R version 3.6.0.

### 2.10. Molecular Dynamics simulations

Two sets of molecular dynamics (MD) simulations were conducted on the VdPelB protein: one in complex with a fully non-methylesterified polygalacturonate decasaccharide, and the other with a fully methylesterified polygalacturonate decasaccharide. Parameters specified by the AMBER14SB_parmbsc1 forcefield [45] were used to create the molecular topologies of the complexes. Each complex was set up in a cubic box with solute-box distances of 1.0 nm and solvated with water molecules specific to the TIP3P water model [46]. Na+ and Cl-ions were added to neutralise the system’s net charge and reach a salt concentration of 0.165 M. Using a steep-descent algorithm with a step size of 0.01, energy minimisation was performed to resolve clashes between particles, with convergence being established at a particle-particle force of 1000 kJ mol-1 nm-1. Particle-particle forces were calculated by considering van der Waals and electrostatic interactions occurring up to 1.0 nm, as well as long-range electrostatics treated in the Fourier space using the Particle Mesh Ewald (PME) summation method. Solvent equilibration was attained post minimisation in two stages: through the nVT and nPT ensembles, to reach constant temperature and pressure, respectively. Equilibration of the solvent under the nVT ensemble was conducted for 1 ns, integrating the equation of motion at a time step of 2 fs. The target reference temperature was 310.15 K, coupled every 0.1 ps using the V-rescale thermostat3. Based on the Maxwell-Boltzmann distribution [47] at 310.15 K, random velocities were then assigned to each particle in the system. Finally, solvent equilibration under the nPT ensemble was conducted for 1 ns, continuing from the last step of the previous equilibration, in terms of particle coordinates and velocities, at a reference temperature of 310.15 K, coupled every 0.1 ps using the V-rescale thermostat [48]. Here, pressure coupling was isotropically coupled every 2.0 ps, at 1 bar, using the Parrinello-Rahman barostat [49]. Particle-particle interactions were computed by constructing pair lists using the Verlet scheme. Short-range van der Waals and electrostatic interactions sampled through a Coulomb potential, were calculated at a cutoff of 1.0 nm. The PME algorithm [50] was used to compute long-range electrostatic interactions beyond this cut-off in the Fourier space, utilising a Fourier grid spacing of 0.16 and a cubic B-spline interpolation level at 4. The simulations were then performed on in-house machines, using GROMACS (Groningen Machine for Chemical Simulations) version 2021.37. Each set of simulations were run for 150 ns each, at a time step of 2 fs, with molecular dynamics trajectories written out every 10 ps. Simulations were replicated 7 times for a total production run time of 1.05 μs per complex. Replicates differed with respect to the random particle velocity sets computed under the nVT ensemble. For analysis, the first 50 ns of each production run were discarded as equilibration time. In-house Python 3 scripts implemented using Jupyter notebooks [51] were used to carry out analyses. Figures were created and rendered with Matplotlib [52] and VMD (Visual Molecular Dynamics)[53].

## 3. Results and Discussion

### 3.1. Sequence and phylogeny analysis

In addition to 9 polygalacturonases (PGs) and 4 pectin methylesterases (PMEs), *V. dahliae* encodes 30 putative endo-pectate lyases (EC 4.2.2.2), exo-pectate lyases (EC 4.2.2.9), endo-pectin lyases (EC 4.2.2.10), belonging to PL1-PL3 and PL9 families, respectively [54]. For hierarchical clustering of Verticilium’s sequences with other PLLs, fifty-one amino acid (aa) sequences encoding putative PLLs, belonging to bacteria, fungi and plants were aligned and a phylogenetic tree was built. Different clades can be distinguished. (**Fig. 1)**. *V. dahliae* PelB (VdPelB, VDAG_04718) clustered with VDAG_05344 (59.68% sequence identity) with close relations to VDAG_05402 (57.19% sequence identity) and VDAG_07566 (56.95% sequence identity). VDAG_05402 and VDAG_05344, that are found in the protein secretome, have orthologs in *V. alfalfa* (VDBG_07839 and VDBG_10041), which were shown to possess putative lyase activity [34, 55]. Plant PLLs from *A. thaliana* (AtPLL21, AtPLL15 and AtPLL18) formed a separate clade with close connections to *A. denitrificans* (AdVexL), a PLL homologue [56]. *D.dadanti* (DdPelI), *Bacillus Sp*. KSM-P15 (BsPel-P15) and *C. bescii* (CbPel1854) formed separate clades similarly to *B.subtilis* and *D. dadantii PLs* (BsPel, DdPelA and DdPelC). The clade corresponding to PNLs consisting of *A. tubingensis* PelA (AtPelA), *A. niger* (AnPelA, AnPnlA and AnPelB), *A*. *luchuensis* AlPelB was closely related to VDAG_07238 and VDAG_08154 which were indeed annotated as putative PNLs [18, 54]. VDAG_06155 was previously named VdPel1 and previously characterized as a pectate lyase [33].

**Fig. 1.**
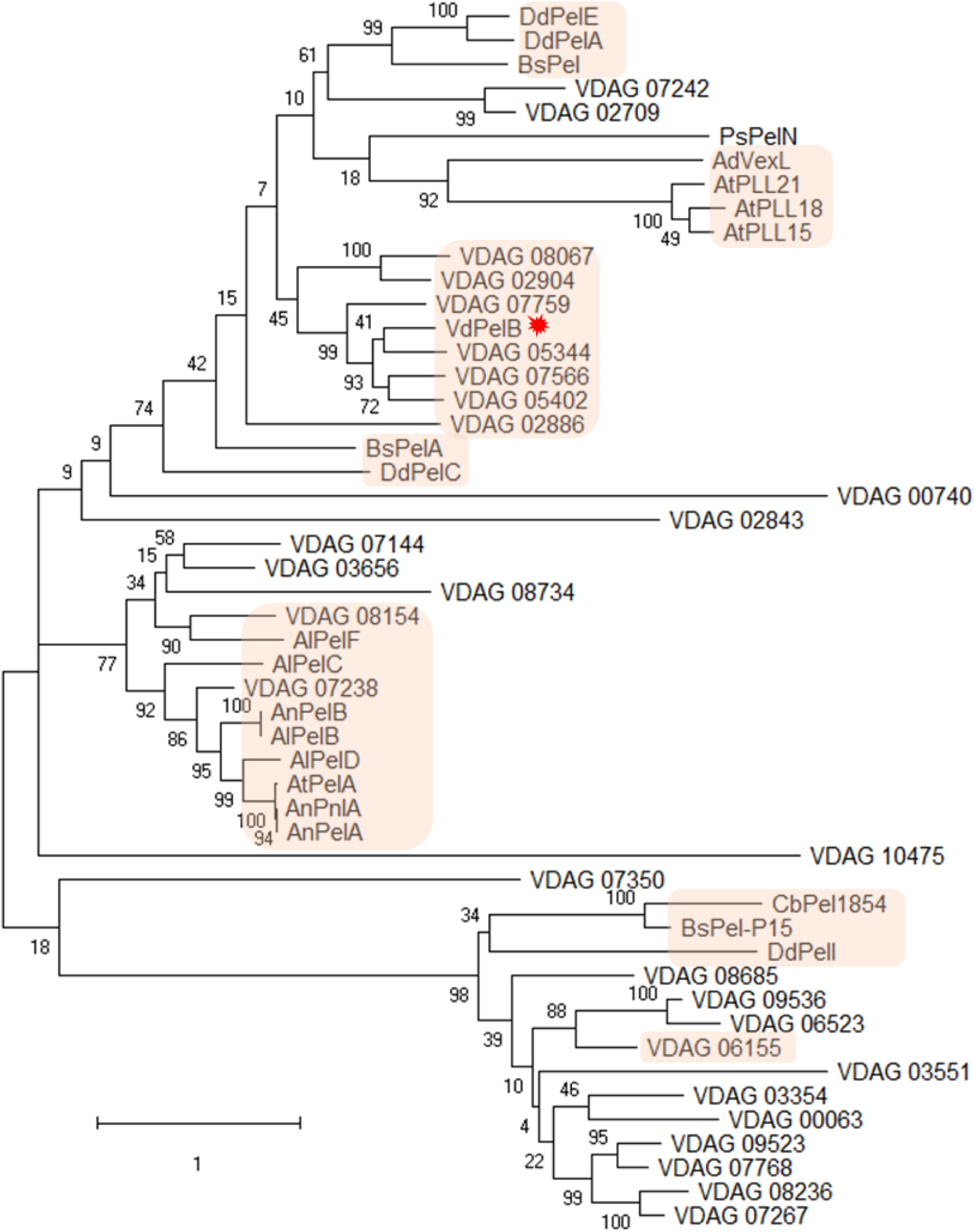
Phylogenetic analysis of *V*. *dahliae* VdPelB with selected PLLs. Phylogentic tree representing *V. dahliae* VdPelB (VDAG_04718, G2X3Y1, red star) amino acid sequence in comparison with PLLs from Verticillium [VDAG_00740 (G2WQU8), VDAG_02904 (G2WXC5), VDAG_05344 (G2X628), VDAG_07242 (G2XBA4), VDAG_07759 (G2XC77), VDAG_02709 (G2WWT0), VDAG_02843 (G2WX64), VDAG_02886 (G2WXA7), VDAG_03656 (G2X1P5), VDAG_05402 (G2X597), VDAG_07144 (G2X9U9), VDAG_07238 (G2XBA0), VDAG_07566 (G2XBY8), VDAG_08067 (G2XD35), VDAG_08154 (G2XDC2), VDAG_08734 (G2XF02), VDAG_10475 (G2XJZ3), VDAG_03354 (G2WZB2), VDAG_03551 (G2WZV9), VDAG_07267 (G2XBC9), VDAG_07768 (G2XC86), VDAG_08236 (G2XDK4), VDAG_06155 (G2X8L4), VDAG_06523 (G2X7R5), VDAG_08685 (G2XEV0), VDAG_09523 (G2XH91), VDAG_09536 (G2XHA4), VDAG_00063 (G2WR80), VDAG_07350 (G2XGG7)], *Arabidopsis thaliana* [AtPLL15 (At5g63180), AtPLL18 (At3g27400), AtPLL21 (At5g48900)], *Dickeya dadanti* [DdPelA (P0C1A2), DdPelC (P11073), DdPelE (P04960), DdPelI (O50325)], *Bacillus subtilis* [BsPel (P39116)], *Bacillus Sp*. KSM-P15 [BsPel-P15 (Q9RHW0)], *Bacillus sp*. N16-5 [BsPelA (D0VP31)], *Aspergillus niger* AnPelA [(Q01172), AnPelB (Q00205), AnPnlA (A2R3I1)], *Achromobacter denitrificans* [AdVexL (A0A160EBC2)], *Aspergillus tubingensis* [AtPelA (A0A100IK89)], *Aspergillus luchuensis* [AlPelB (G7Y0I4)], *Acidovorax citrulli* [AcPel343, (A1TSQ3)], *Paenibacillus sp.* 0602 [PsPelN (W8CR80)], *Caldicellulosiruptor bescii* [CbPel1854 (B9MKT4)]. Maximum likelihood tree was constructed with 1000 bootstrap replicates. Most important clades are indicated in orange squares while VdPelB is marked by red star. Amino acids sequences were retrieved from Uniprot and TAIR.

### 3.2. Cloning, expression and purification of VdPelB

The *VdPelB* (*VDAG_04718*) gene consists of 1 single exon of 1002 bp length. The coding sequence, minus the putative signal peptide was PCR-amplified using gene-specific primers, ligated in pPICZαB vector and expressed in *P. pastoris* heterologous expression system, allowing its secretion in the culture media. Secreted VdPelB consisted of 343 aa, including the poly-histidine tag at the C-terminus used for affinity chromatography purification. After purification, VdPelB had an apparent molecular mass of ∼38 kDa (**Fig. S1A**), higher than what was predicted on the basis of the amino acid sequence (33.8 kDa). However, this shift is likely to correspond to the tags (His and C-myc) and to the presence of 19 putative O-glycosylation sites, as predicted by NetOGlyc 4.0 Server.

### 3.3. VdPelB has a right-handed parallel β-helix fold

VdPelB was crystallized and its 3D structure was determined using X-ray diffraction. VdPelB crystallized in monoclinic P 1 2_1_ 1 asymmetric unit. Four data sets, collected from the same crystal at 1.2 Å resolution, were integrated, scaled and merged. There are two molecules in the asymmetric units: chains A and B are highly similar with a Cα root mean square deviation (rmsd) value of 0.227 Å (**Fig. S2A**). The VdPelB structure consists of 298 amino acids (aa) with 18 aa at the N-terminus and 27 aa at the C-terminus that were not resolved because of poor electron densities, while overall electron densities were well defined. While no N-glycosylation sites could be revealed on the VdPelB structure, six O-glycosylation sites carrying mannose are visible for each molecule: T22, T44, T45, T46, S48 and T54, in accordance with the shift in size previously observed (**Fig. S2A**). During data acquisition no heating of the crystal was observed, as shown by low B factors and good occupancies (**Fig. S3A and B**). The final models’ geometry, processing and refinement statistics are summarized (**Table 1**). VdPelB’s structure has been deposited in the Protein Data Bank as entry 7BBV.

**Table 1.** Data collection, processing and refinement for VdPelB.

| Characteristics | VdPeIB |
| --- | --- |
| Data collection |  |
| Diffraction source | PROXIMA2 |
| Wavelength (Å) | 0.980 |
| Temperature (°C) | 100.15 |
| Detector | DECTRIS EIGER X 9M |
| Crystal-to-detector distance (mm) | 115.02 |
| Rotation range per image (°) | 0.1 |
| Total rotation range (°) | 360 |
| Crystal data |  |
| Space group | P1 21 1 |
| *a*, *b*, *c* (Å) | 60.89, 59.69, 93.87 |
| $\alpha$, $\beta$, $\gamma$, (°) | 90.00, 96.35, 90.00 |
| Subunits per asymmetric unit | 2 |
| Data statistics |  |
| Resolution range (Å) | 60.52-1.2 (1.243 - 1.2) |
| Total No. of reflection | 8536161 (112943) |
| No. of unique reflection | 207497 (19983) |
| No. of reflections, test set | 10362 (1003) |
| $R_{\text{merge}}$ (%) | 0.1036 (4.709) |
| Completeness (%) | 99.30 (96.11) |
| $\langle I/\sigma(I) \rangle$ | 18.03 (0.22) |
| Multiplicity | 41.1 (5.6) |
| $CC_{1/2}$ (%) | 1 (0.116) |
| Refinement |  |
| $R_{\text{crys}}/R_{\text{free}}$ (%) | 17.3 / 19.7 |
| Average B – factor (Å<sup>2</sup>) | 33.18 |
| No. of non-H atoms |  |
| Protein | 9097 |
| Ion | 2 |
| Ligand | 234 |
| Water | 1052 |
| Total | 10385 |
| R.m.s. deviations |  |
| Bonds (Å) | 0.013 |
| Angles (°) | 1.39 |
| Ramachandran plot |  |
| Most favoured (%) | 95.1 |
| Allowed (%) | 4.90 |
| Outlier (%) | - |

VdPelB has a right-handed parallel β-helix fold which is common in pectinases [57]. The β-helix is formed by three parallel β-sheets - PB1, PB2 and PB3 which contain 7, 10 and 8 β-strands, respectively. Turns connecting the PB1-PB2, PB2-PB3 and PB3-PB1 β-sheets are named T1-turns, T2-turns and T3-turns, respectively, according to Yoder and Jurnak (**Fig. 2A, S4A and B)** [58]. T1-turns consist of 2-14 aa and builds the loop around the active site on the C-terminus. T2 turns mostly consist of 2 aa with Asn being one of the predominant aa, forming an N-ladder with the exception of an N245T mutation in VdPelB [15, 20]. VdPelB has a α-helix on N-terminus end that shields the hydrophobic core and is commonly conserved in PLs and PGs [15, 59], while the C–terminus end is also protected by tail-like structure carrying one α-helix. Interestingly N- and C-terminus tails pack against PB2 (**Fig. 2A**). There are only two Cys (C25 and C137) that do not form a disulphide bridge.

**Fig. 2.**
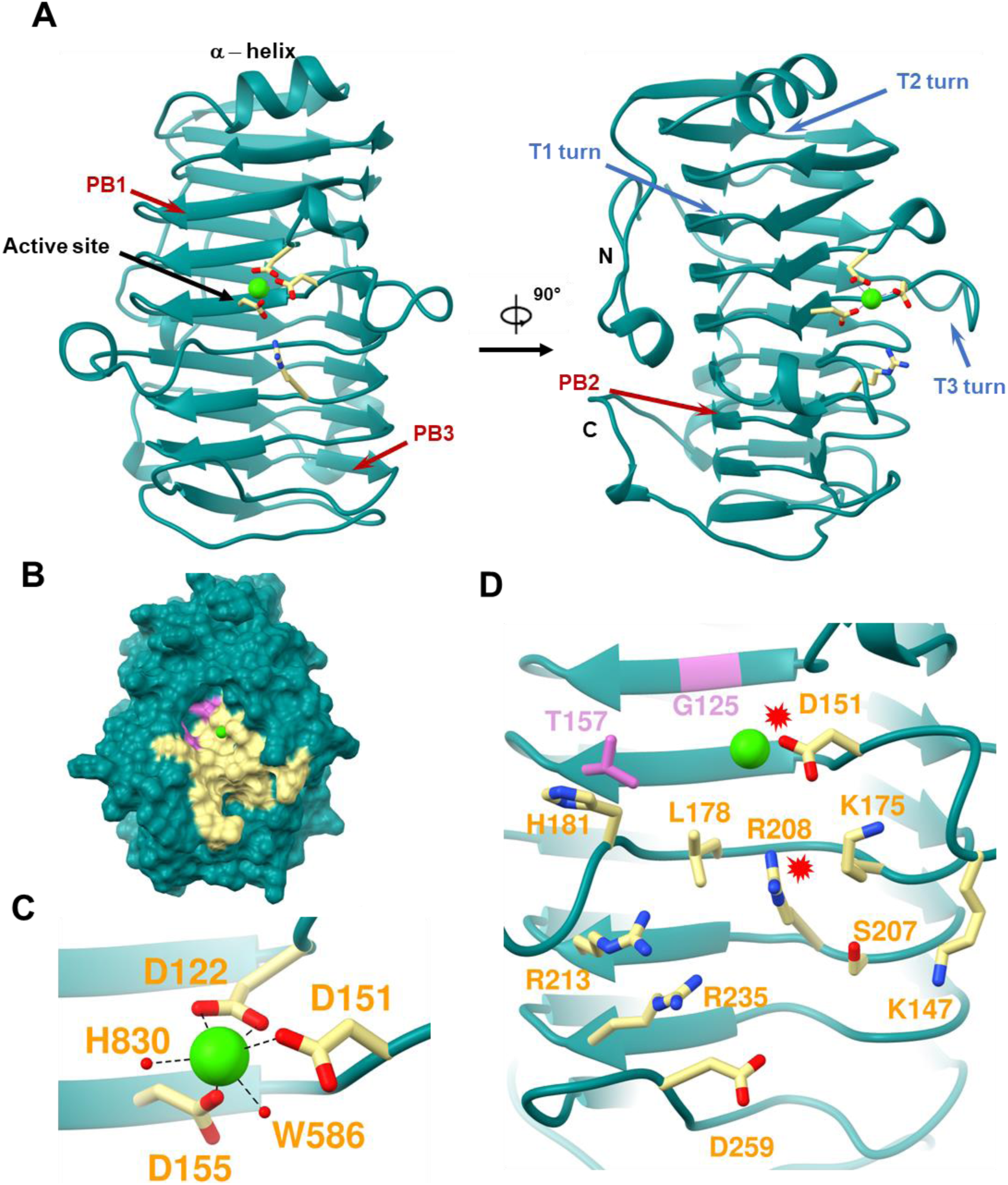
Structure determination of VdPelB. A) Ribbon diagram of VdPelB crystalized in P1 21 1 space group. VdPelB is a right-handed parallel β helical structure consisting of β strands (red arrows) and turns (blue arrows). VdPelB active site’s aa are yellow-colored while Ca atom is green. B) Surface representation of VdPelB binding groove. C) Active site of VdPelB highlighting conserved aa and atoms interacting with Ca. D) Structure of VdPelB binding groove highlighting aa involved in the interaction (yellow) and aa not of previously characterized in PLs (plum). Red stars indicates aa from the active site.

Sequence and structural alignments show that VdPelB belong to the PL1 family. VdPelB shares the highest structural similarity with BsPelA (PDB: 3VMV), with 30.06% sequence identity and Cα rmsd of 1.202 Å. The second-best structural alignment was with DdPelC (PDB: 1AIR) with 24.20% identity and Cα rmsd of 1.453 Å [41, 60]. Both of these structures lack the long T3 loop described in *A.niger* pectate lyase (AnPelA, PDB: 1IDJ**, Fig. S2B** and **S5**) [15]. The putative active site is positioned between the T3 and T2 loops (**Fig. 2A and B**).

### 3.4. Active site harbours Ca^2+^ that is involved in catalysis

The VdPelB active site is well conserved, harbouring strictly conserved acidic and basic aa that are required for Ca^2+^ binding. Previously reported structures showed that two Asp (D122 and D155) and one Arg (R208) in VdPelB, are conserved, while D151 can be mutated to Glu, or Arg in PNLs (**Fig. 2C** and **D**, Mayans et al., 1997). Other conserved aa in VdPelB include K175 and R213, with K175 being responsible for binding the carboxyl oxygen while R213 hydrogen bonds to C-2 and C-3 of GalA (**Fig. 2D**) [29, 61]. Mutating any of these aa leads to decreased enzyme activity [62]. In VdPelB, Ca^2+^ ion is directly coordinated by D122, two carboxyl oxygen, D151, D155 and two water molecules (W568 and W830, **Fig. 2C**). In addition, mutation of D122T (VdPelB numbering) in BsPelA, is responsible for reduced affinity for Ca^2+^ [41]. In the catalytic mechanism, Ca^2+^ is directly involved in acidification of the proton absorption from C5 and elimination of group from C4, generating an unsaturated product. R208 act as a base, similarly to the hydrolysis in the reaction mechanism of the GH28 family [63, 64].

### 3.5. Structural analysis of VdPelB suggests a PL activity mediated by peculiar specificities

The VdPelB binding groove comprises a number of basic and acidic aa including K147, D151, K175 L178, H181, S207, R208, R213 and R235 and D259 (**Fig. 2C and D**), that have previously been shown to be characteristics of PLs. This would suggest an enzyme activity on low DM pectins as aa positioned at the binding groove were indeed shown to differ between PNL and PL [15,21,24,29]. These aa are indeed mutated in Arg, Trp, Tyr, Gln and Gly in PNLs, which, by reducing their charge, would favour higher affinity for highly methylesterified pectins [15, 29]. In VdPelB, as the T3 loop is missing, there are no equivalent to PNL specific W66, W81, W85, W151 aa (**Fig. S2B,** AnPelA numbering) and, moreover, W212 and W215 are replaced by K175 and L178 in VdPelB. In BsPelA and DdPelC, these aa are replaced by K177/K190 and L180/L193, highlighting the high conservation of amino acids in VdPelB/BsPelA/DdPelC and subsequently in PLs. When a DP4 ligand from DdPelC crystal structure is superimposed to VdPelB, hydrogen bonds and van der Waals interactions are visible with the above-mentioned aa (**Fig. S6)** [15,21,29,41].

Interestingly, despite this rather conserved PL-related binding grove, VdPelB harbors, in the vicinity of the active site, G125 and T157 that are not present in the well characterized DdPelC [60]. At these positions, DdPelC, which was shown to be a *bona fide* PL with a high activity on polygalacturonic acid and alkaline pH with Ca^2+^-dependency, harbours Arg and Lys [15,61,65]. In that respect, the presence of G125 and T157 in VdPelB is similar to that identified in *Bacillus sp.* Pel-22, Pel-66, and Pel-90 and *Bacillus sp.* which showed activity on both PGA and high methylesterified pectins (**Fig. 2B** and **D**) [22,66,67]. The DdPelC aa being overall positively charged, could explain the binding preference towards non-methylesterified, negatively charged substrates in the vicinity of the active site. In contrast, considering the size of Gly and Thr, they would sterically accommodate the increased size of methylesterified substrate. G125 and T157 could therefore account for a potential dual activity of VdPelB. Moreover, in previously characterized PLs and PNLs there is the presence of a small aa, Ser or Ala that replace H181. While H181 interacts directly with the substrate these aa do no provide this interaction instead the primary Ca^2+^ in the active site induce a substrate conformation that could be recognized by PLs [41]. Finally, L178 is positioned in-between the catalytic Ca^2+^ and R208 and is involved in substrate binding making it a perfect candidate to assess its importance (**Fig. 2D**).

### 3.6. Molecular dynamic simulations show higher dynamics of VdPelB in complex with methylesterified substrates

To determine how the structure of VdPelB and the observed differences in amino-acidic composition might influence the affinity with differently methylated substrates, we performed MD simulations on VdPelB in complex with either a non-methylesterified or fully methylesterified decasaccharides, which are able to occupy the entire binding groove (**Fig. 3**). MD simulations show substantially differential dynamic profiles for oligosaccharides with and without methylesterification. Expectedly, the dynamics is lowest in proximity of the catalytic subsite (+1) and increases consistently towards both the reducing and non-reducing ends of the substrate: with the highest dynamics found at the non-reducing end (**Fig. 3A**).

**Fig. 3.**
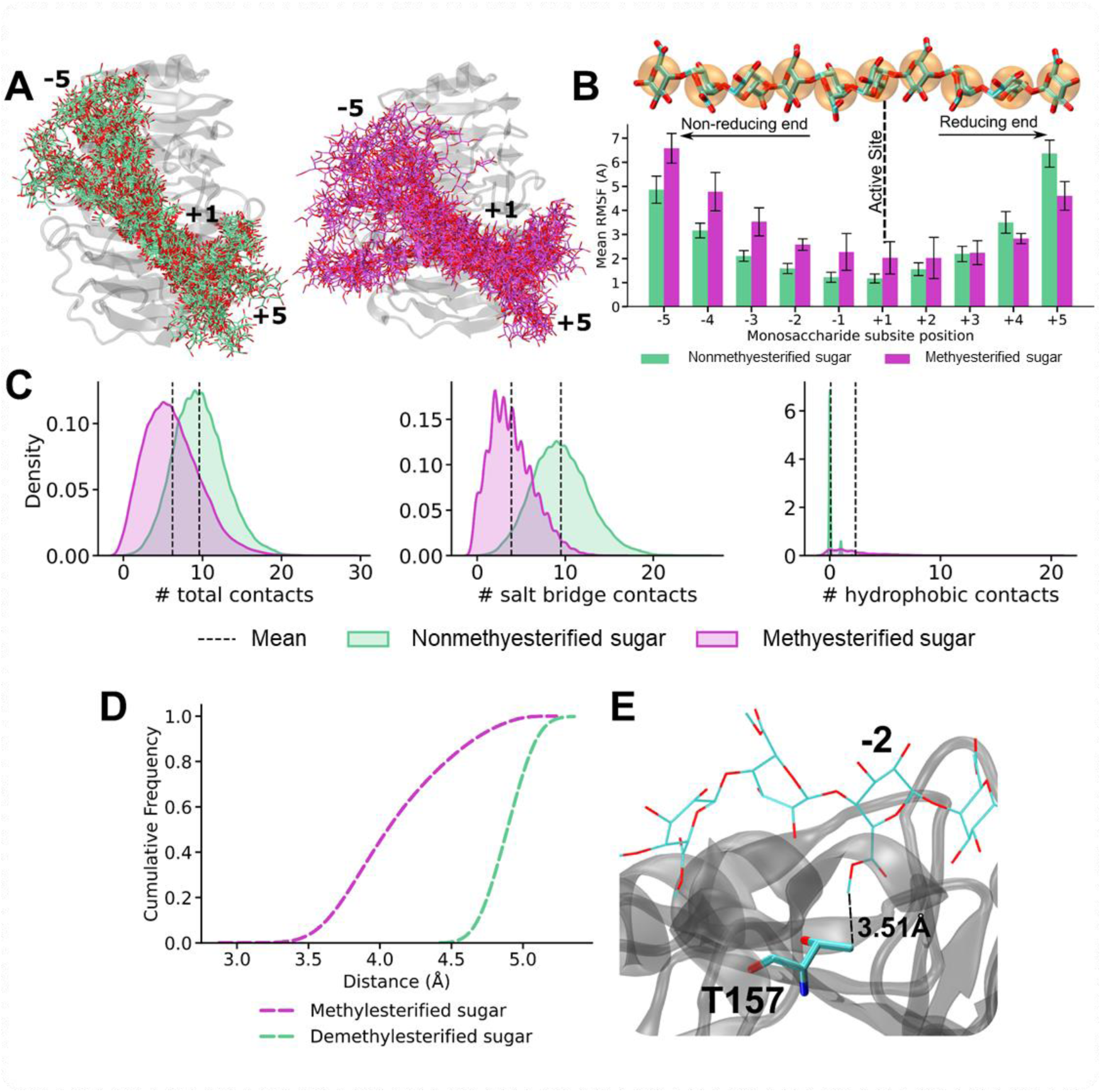
VdPelB substrate dynamics in complex with fully de-methylesterified and methylesterified complex. A) Ensembles of non-methylesterified (green, left panel) and methylesterified (pink, right panel) HG decasaccharide at every 100 frames for the simulated VdPelB complexes. Substrate within the enzymes’ binding grooves are labelled from −5 (HG-non-reducing end) to +5 (HG-reducing end) B) RMSF of non-methylesterified (green) and methylesterified (pink) HG bound across the binding groove of VdPelB. Numbers indicate the beta sheet position form the active site. C) Analysis of the contacts between VdPelB and non-methylesterified (green) and methylesterified substrate (pink). D) Distance between T157K residue and the substrate residues plotted as cumulative frequency. E) T157 hydrophobic interaction with methyl-ester of methylesterified substrate.

A quantitative estimation of the substrate dynamics was obtained by monitoring the root square mean fluctuations (RMSF) of each sugar residue and shows that a polygalacturonate substrate associates more stably in the subsites of the binding groove placed towards the sugar’s reducing end (subsites −1 to −5), with differences between methylesterified and non-methylesterified substrates approaching the obtained standard deviation. In some cases, the dynamics reversed for non-methylesterified sugars, with a higher RMSF for de-methylesterified sugars towards the saccharide’s reducing end (**Fig. 3B**). Moreover, and in line with an observed positively charged binding groove, non-methylesterified substrates retain a higher number of contacts than methylesterified sugars, with salt-bridges contributing the most to the observed differences (**Fig. 3C**).

We then additionally and specifically focused on the analysis of the interactions made by T157, which is nested with the −2 subsite of the binding groove. Simulations sample consistently higher contacts formation between a methylesterified sugar docked in subsite −2 and T157, with distances shifted to lower values when compared to a non-methylesterified sugar (**Fig. 3D**), even the RMSF of non-methylesterified monosaccharides docked in subsite −2 experience on average, significantly lower dynamics. Altogether, the formation of a larger number of contacts between methylesterified saccharides docked in the −2 subsite suggests and active role of T157 in the binding of methylesterified chains: with the butanoic moiety of T157 engaging in hydrophobic interactions with the methyl-ester presented by methylesterified sugar units **(Fig. 3E**). While T157 is seen to have an active role in engaging with the substrate’s hydrophobic moieties, an active role of G125, also in the same subsite, was not observed but it is plausible that the minimal size of G125 would increase the accommodability of methylesterified sugars in that position.

### 3.7. Biochemical characterization of VdPelB

We first determined the activity of VdPelB by following the release of 4,5-unsaturated bonds which can be detected at 235 nm using UV spectrophotometer. VdPelB activity was first tested for its dependency towards Ca^2+^. Using a standard PL assay, with PGA as a substrate, an increase in the VdPelB activity was measured in presence of calcium. In contrast in presence of EDTA, used as a chelating agent, no activity was detected, confirming the calcium-dependent activity of the enzyme **(Fig. S7A)** [15]. Activity measured in absence of added CaCl_2_ reflects the presence of calcium from the culture media that is bound to VdPelB during production, and previously identified in the 3D structure. We tested the effects of increasing CaCl_2_ concentrations and showed that the maximum activity was already reached when using as low as 0.125 µM (**Fig. S7B**). To test the substrate-dependence of VdPelB, four substrates of increasing degrees of methyl-esterification were used. VdPelB showed the highest activity on PGA, with less than 10% of the maximum activity measured on the three others substrates (**Fig. 4A**). This shows that, as inferred from above-mentioned structural and dynamical data, VdPelB act mainly as a PL although it can still show residual activity on high DM pectins. Considering this, PGA was used as substrate to test the pH-dependence of the enzyme’s activity in sodium acetate and Tris-HCl buffers (**Fig. 4A**). VdPelB was most active at pH 8, with only a slight decrease in activity at pH 9 (93%). In contrast, the relative activity at pH 5-7 was close to null. The pH optimum for VdPelB was the same as *B. fuckeliana* Pel (pH 8) [68], close to that reported for *D. dadantii* PelN (pH 7.4) [69], but was lower to that measured for *B.clausii* Pel (pH 10.5) [70]. In contrast, the pH optimum was higher compared to five PNLs from *Aspergillus* sp. AaPelA (pH 6.1), AtPelA (pH 4.5), AtPelA (pH 6.4), AtPelD (pH 4.3) [18] and *A. parasiticus* Pel (pH 4) [19]. The optimum temperature assay showed that VdPelB was most active at 35°C (**Fig. S8**). VdPelB appeared less heat-tolerant as compared to thermophilic PLLs reported from Bacillus sp. RN1 90°C [71], *B. clausii* Pel, 70°C [70], *B. subtilis* Pel168, 50°C [72]. However, its optimum temperature is in the range of that measured for *X. campestris* Pel [73] and cold-active Pel1 from *M. eurypsychrophila* [74]. The lack of disulphide bridges previously shown in the structure could be responsible for the lower stability of the enzyme at high temperatures, in comparison with previously characterized PLs [75].

**Fig. 4.**
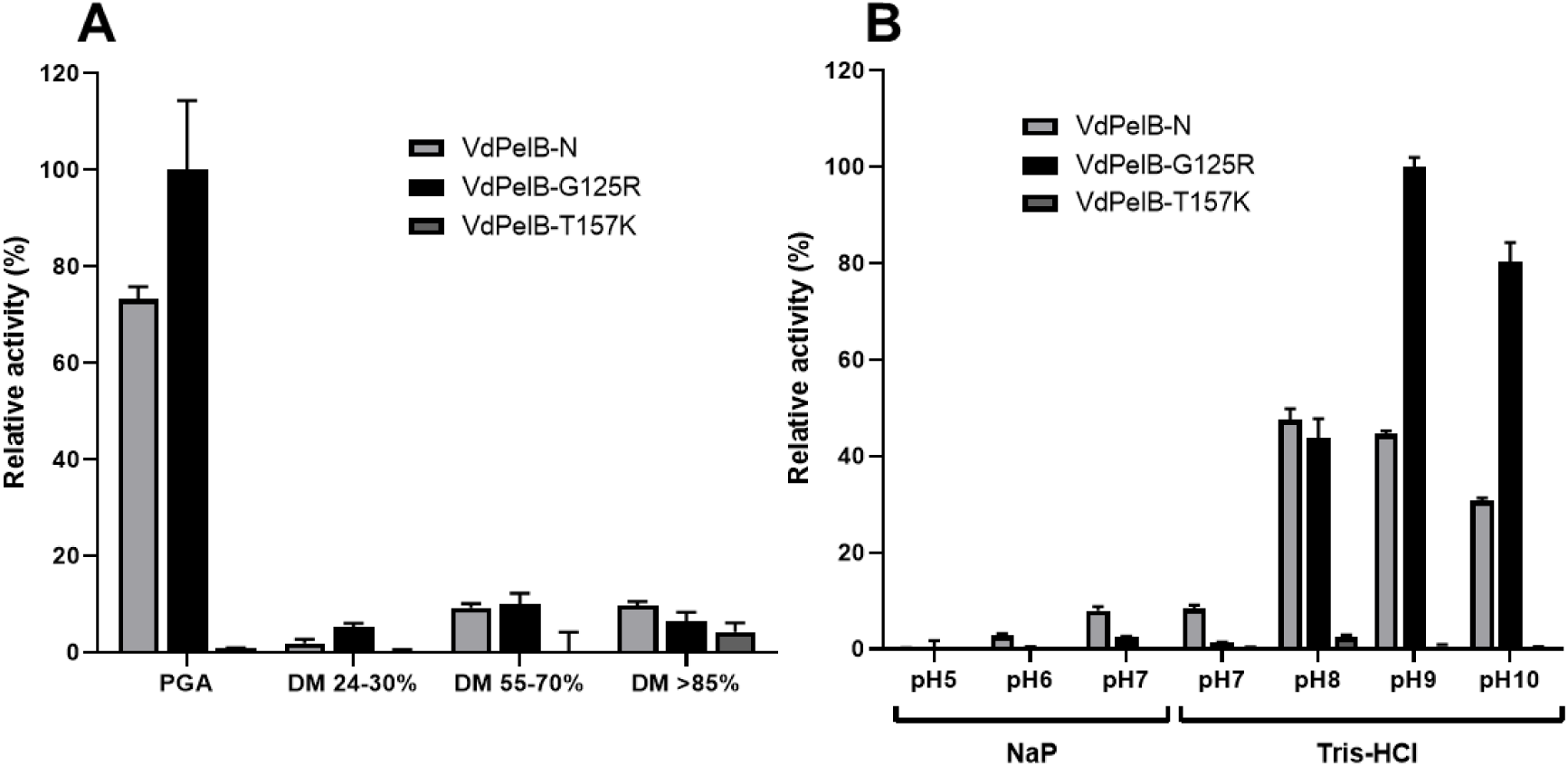
Biochemical characterization of VdPelB. A) Substrate-dependence of VdPelB-N, G125R and T157K. The activities were measured after 12 min of incubation with PGA, pectins DM 24-30%, DM 55-70%, DM>85% with addition of Ca^2+^ at 35°C. B) pH-dependence of VdPelB-N, G125R and T157K activity. The activities were measured after 12 min of incubation with PGA in sodium phosphate (NaP) and Tris-HCl buffer at 35°C. Values correspond to means ± SE of three replicates.

### 3.8. Mutation of specific amino-acids affects VdPelB activity

Considering the fine structure of VdPelB structural we generated mutated forms of the enzymes for five amino acids that likely to be involved in the catalytic mechanism and/or substrate binding: G125R, D151R, T157K, L178K and H181A. Enzymes were produced in *P. pastoris* and purified (**Fig. S1B**). If the importance of some of these aa (i.e. D151 and H181) in the catalytic mechanism was previously shown for others PL [41, 76], our study highlights the key role of some novel aa in the catalytic mechanism. The activities of mutants were tested at iso-quantities of wild-type enzyme. Surprisingly, the G125R mutant was 25% more active on PGA compared to the native enzyme and a shift in the optimum pH was observed (**Fig. 4A**). While native enzyme was most active at pH8 with slight decrease in activity at pH9, the activity of G125R mutant was increased by circa 50% at pH9 and pH10 (**Fig. 4B**). Both enzymes kept the same activity at pH8. The substitution of Gly with Arg, present in a number of previously characterized PL could facilitate the interaction with the substrate and basification of the active site thanks to its physio-chemical properties [60]. The activity of T157K was closed to null when tested on PGA and different pHs. The introduction of a Lys, a chemically different and larger aa is likely to introduce steric clashes notably with H181 and L178 that are important for the activity. The H181A mutant lost much of its activity as there are less interaction/recognition with the substrate. No activity for D151R and L178K mutants were observed in line with the fact that D151 is an active site aa that binds the Ca^2+^. Mutation of this amino-acid is negatively impairing the functioning of the enzyme (**Fig. S7A** and **B)** [41, 76]. We can hypothesize that L178K mutation positioned in between Ca^2+^, R208 and the substrate, induces specific substrate conformation that diminishes the direct interaction between the enzyme catalytic centre and the substrate, which translates to loss of activity (**Fig. S9**).

### 3.9. Identification of the OGs released by VdPelB from commercial and cell wall pectins

To further understand the specificity of VdPelB on different substrates, we performed LC-ESI-MS/MS to determine the profiles of digestion products (OGs) and to compare with that of commercially available *Aspergillus sp.* Pel (AsPel, **Fig. S10**). To be fully comparable, digestions were realized, for each substrate, at iso-activities for the two enzymes. On the basis of digestion profiles, we identified 48 OGs and created a dedicated library that was used for identification and integration of peaks (**Table 2**). MS^2^ fragmentation allowed determining the structure of some of the OGs (**Fig. S11** and **S12)**. The OGs released by either of the enzymes were mainly corresponded to 4,5-unsaturated OGs, which is in accordance with β-eliminating action of PLLs. When using pectins DM 20-34% and at low enzyme’s concentration (VdPelB-2), VdPelB mainly released non-methylesterified OGs of high DP (GalA_5_, GalA_6_, GalA_7_, GalA_8_, GalA_9_, GalA_10_) that were subsequently hydrolysed when using more concentrated VdPelB (VdPelB-1, **Fig. 5A**). These digestion products strikingly differed to that generated by AsPel, that are methylesterified OGs of higher DP (GalA_4_Me_1_, GalA_4_Me_2_, GalA_5_Me_1_, GalA_6_Me_2_, GalA_11_Me_3_…), thus showing distinct enzymatic specificities. Altogether, these first results unequivocally shows VdPelB act as an endo-PL, and that its processivity differ to that of AsPel, which act as well as an endo-PL [9]. When using sugar beet pectins, that are known to be highly acetylated (DM 42%, DA 31%), VdPelB released acetylated OGs (GalA_2_Ac_2_, GalA_3_Ac_1_, GalA_4_Ac_1_, GalA_5_Ac_1_), while AsPel showed much lower activity and relative abundance of these OGs (**Fig. 5B**). Previous reports have shown that differences exist between PNL, in particular with regards to acetyl substitutions [18]. In contrast, AsPeI released mainly methylesterifed OGs (GalA_6_Me_1_, GalA_8_Me_1_, GalA_10_Me_3_…).

**Table 2.** List of oligogalacturonides identified by LC-MS/MS analysis. For each OG, elemental composition, retention time (RT) and ion mass used for the analysis are highlighted. In the case of detection of mono and di-charged OGs, the italicized and **BOLD** mass, corresponding to the more intense ion, were used for the quantification. * indicates non-unsaturated OGs.

| Name | Elemental composition | RT (min) | [M-H]- | [M-2H]2- |
| --- | --- | --- | --- | --- |
| GalA1* | C6H10O7 | 9.3 | 193.0353763 |  |
| GalA1Ac2 | C10H12O8 | 9.1 | 259.045941 |  |
| GalA2* | C12H18O13 | 8.88 | 369.0674644 |  |
| GalA2 | C12H16O12 | 8.95 | 351.0568997 |  |
| GalA2Ac2 | C16H20O14 | 8.67 | 435.0780291 |  |
| GalA2Me1* | C13H20O13 | 8.85 | 383.0831145 |  |
| GalA2Me1 | C13H18O12 | 8.92 | 365.0725498 |  |
| GalA3* | C18H26O19 | 8.56 | 545.0995525 |  |
| GalA3 | C18H24O18 | 8.61 | 527.0889878 |  |
| GalA3Ac1 | C20H26O19 | 8.59 | 569.0995525 |  |
| GalA3Me1 | C19H26O18 | 8.56 | 541.1046379 |  |
| GalA3Me2* | C20H30O19 | 8.62 | 573.1308527 |  |
| GalA4* | C24H34O25 | 8.27 | 721.1316406 |  |
| GalA4 | C24H32O24 | 8.33 | **703.1210759** | 351.0568997 |
| GalA4Ac1 | C26H34O25 | 8.31 | 745.1316406 |  |
| GalA4Me1* | C25H36O25 | 8.37 | 735.1472907 |  |
| GalA4Me1 | C25H34O24 | 8.32 | **717.136726** | 358.0647248 |
| GalA4Me2 | C26H36O24 | 8.3 | 731.1523761 |  |
| GalA5* | C30H42O31 | 8.04 | 897.1637287 |  |
| GalA5 | C30H40O30 | 8.09 | **879.153164** | 439.0729438 |
| GalA5Ac1 | C32H42O31 | 8.09 | 921.1637287 |  |
| GalA5Me1* | C31H44O31 | 8.11 | **911.1793788** | 455.0860512 |
| GalA5Me1 | C31H42O30 | 8.13 | **893.1688141** | 446.0807688 |
| GalA5Me2* | C32H46O31 | 8.13 | 925.1950289 |  |
| GalA5Me2 | C32H44O30 | 8.03 | 907.1844642 | **453.0885939** |
| GalA6* | C36H50O37 | 7.84 |  | 536.0942702 |
| GalA6 | C36H48O36 | 7.82 | 1055.185252 | **527.0889878** |
| GalA6Me1 | C37H50O36 | 7.92 | 1069.200902 | **534.0968129** |
| GalA6Me2 | C38H52O36 | 7.83 | 1083.216552 | **541.1046379** |
| GalA7* | C42H58O43 | 7.65 |  | 624.1103142 |
| GalA7 | C42H56O42 | 7.62 | 1231.21734 | **615.1050318** |
| GalA7Me1 | C43H58O42 | 7.72 | 1245.23299 | **622.1128569** |
| GalA7Me2* | C44H62O43 | 7.75 |  | 638.1259643 |
| GalA7Me2 | C44H60O42 | 7.72 | 1259.24864 | **629.1206819** |
| GalA8 | C48H64O48 | 7.45 |  | 703.1210759 |
| GalA8Me1* | C49H68O49 | 7.53 |  | 719.1341833 |
| GalA8Me1 | C49H66O48 | 7.52 | 1421.265078 | **710.1289009** |
| GalA8Me2* | C50H70O49 | 7.55 |  | 726.1420083 |
| GalA8Me2 | C50H68O48 | 7.58 | 1435.280728 | **717.136726** |
| GalA9* | C54H74O55 | 7.29 |  | 800.1424023 |
| GalA9 | C54H72O54 | 7.3 |  | 791.1371199 |
| GalA9Me1 | C55H74O54 | 7.34 |  | 798.144945 |
| GalA9Me2 | C56H76O54 | 7.37 |  | 805.15277 |
| GalA9Me2 | C56H76O54 | 7.41 |  | 805.15277 |
| GalA9Me3 | C57H78O54 | 7.45 |  | 812.1605951 |
| GalA10 | C60H80O60 | 7.15 |  | 879.153164 |
| GalA10Ac1 | C62H82O61 | 7.29 |  | 900.1584463 |
| GalA10Me1 | C61H82O60 | 7.19 |  | 886.160989 |
| GalA10Me2* | C62H86O61 | 7.24 |  | 902.1740964 |
| GalA10Me2 | C62H84O60 | 7.24 |  | 893.1688141 |
| GalA10Me3 | C63H86O60 | 7.3 |  | 900.1766391 |
| GalA11 | C66H88O66 | 7.03 |  | 967.169208 |
| GalA11Ac1 | C68H90O67 | 7.14 |  | 988.1744904 |
| GalA11Me1 | C67H90O66 | 7.06 |  | 974.1770331 |
| GalA11Me2 | C68H92O66 | 7.09 |  | 981.1848581 |
| GalA11Me3 | C69H94O66 | 7.14 |  | 988.1926832 |
| GalA12Me2 | C74H100O72 | 6.96 |  | 1069.200902 |
| GalA12Me3 | C75H102O72 | 6.99 |  | 1076.208727 |
| GalA12Me4 | C76H104O72 | 7.0419 |  | 1083.216552 |
| GalA13Me2 | C80H108O78 | 6.82 |  | 1157.216946 |
| GalA13Me3 | C81H110O78 | 6.86 |  | 1164.224771 |
| GalA13Me4 | C82H112O78 | 6.9 |  | 1171.232596 |
| GalA14Me3 | C87H118O84 | 6.73 |  | 1252.240815 |
| GalA14Me4 | C88H120O84 | 6.77 |  | 1259.24864 |
| GalA15Me5 | C95H130O90 | 6.69 |  | 1354.272509 |

**Fig. 5.**
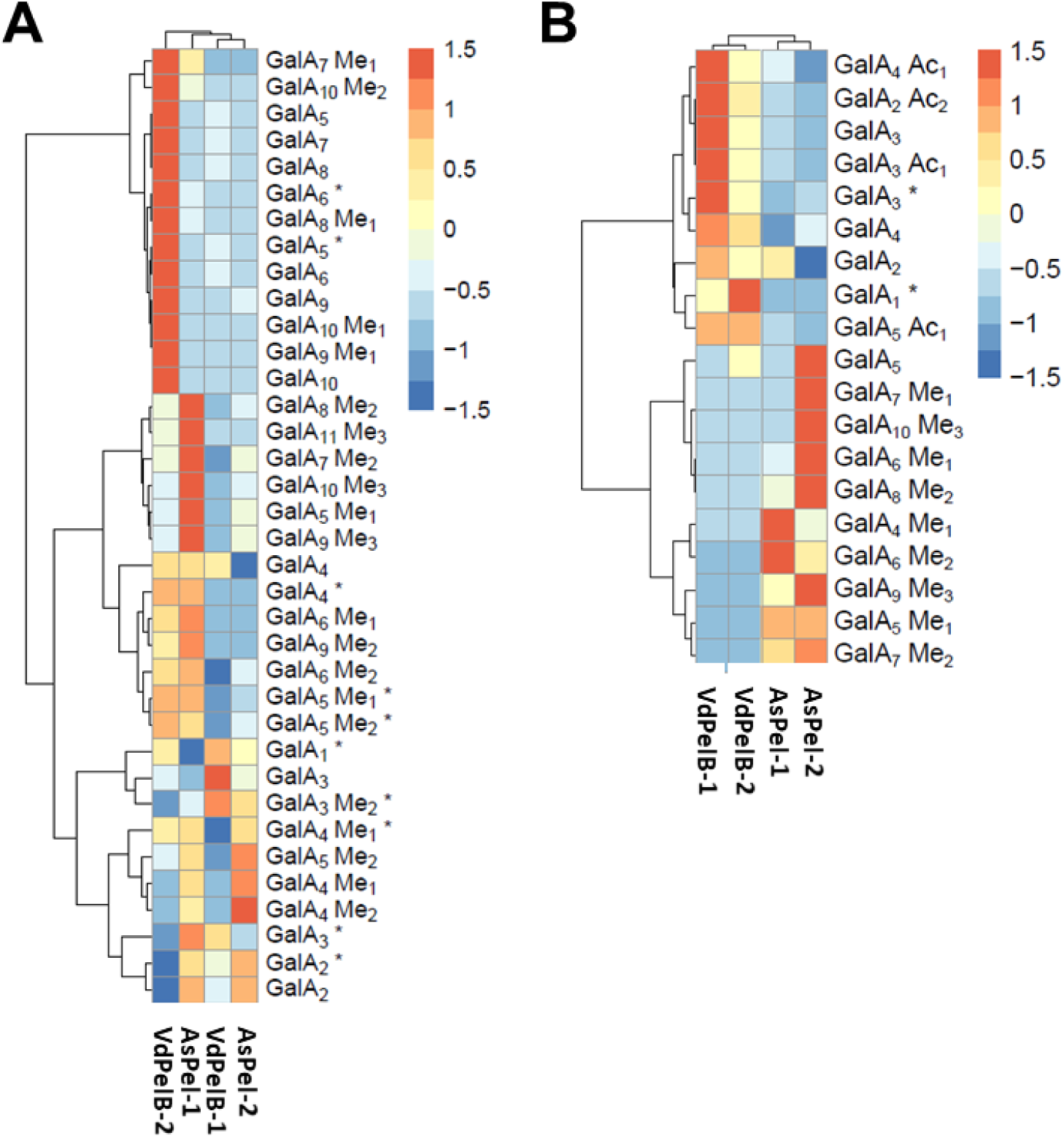
Analysis of OGs produced by the action of VdPelB and AsPeI on pectins of various degrees of methylesterification and acetylation. OGs were separated by SEC and analysed by MS/MS. A) Pectins DM 24-30%. B) Sugar beet pectins (DM 42% DA 31%). Substrates were digested overnight at 40°C and pH 8 using isoactivities of VdPelB and AsPel. Enzyme concentrations are stated in Table S2. Subscript numbers indicate the DP, DM and DA. * indicate non-unsaturated OGs. Values correspond to means of three replicates.

*V. dahliae* and closely related fungus from the same genus are known flax pathogens, where they use their enzymatic arsenal, that includes pectin degrading enzymes, for penetrating the host cell leading to infection [30]. To assess the potential role of VdPelB in flax pathogenicity we digested root cell walls from two flax cultivars, Evea (Verticillium-partially resistant), and Violin (Verticillium-susceptible), and we compared the OGs released. On root cell walls, VdPelB released mainly unsaturated OGs up to DP5 (**Fig. 6**). From similar starting root material, the OG total peak area detected was five times lower for Evea compared to Violin, suggesting that it is less susceptible to digestion by VdPelB. OGs released by VdPelB were mainly non-methylesterified but could be acetylated (GalA_2_, GalA_3_, GalA_3_Ac_1_, GalA_4_, GalA_4_Ac_1_ GalA_5_, GalA_5_Ac_1_), and the abundance of GalA_3_ and GalA_4_ was five and fifteen times higher in Violin, respectively. These data together with that obtained from sugar beet pectins strongly suggests that VdPelB preference is for non-methylesterified and acetylated substrates. Our data suggest that cell wall structure differ between the two cultivars and that VdPelB could determine Verticillium pathogenicity thanks to a better degradation of the cell wall pectins of sensitive cultivars [35]. Similarly, VdPel1 was previously identified as virulence factor, where the deletion of this gene decreased virulence in tobacco, as compared with the wild-type Verticillium [33].

**Fig. 6.**
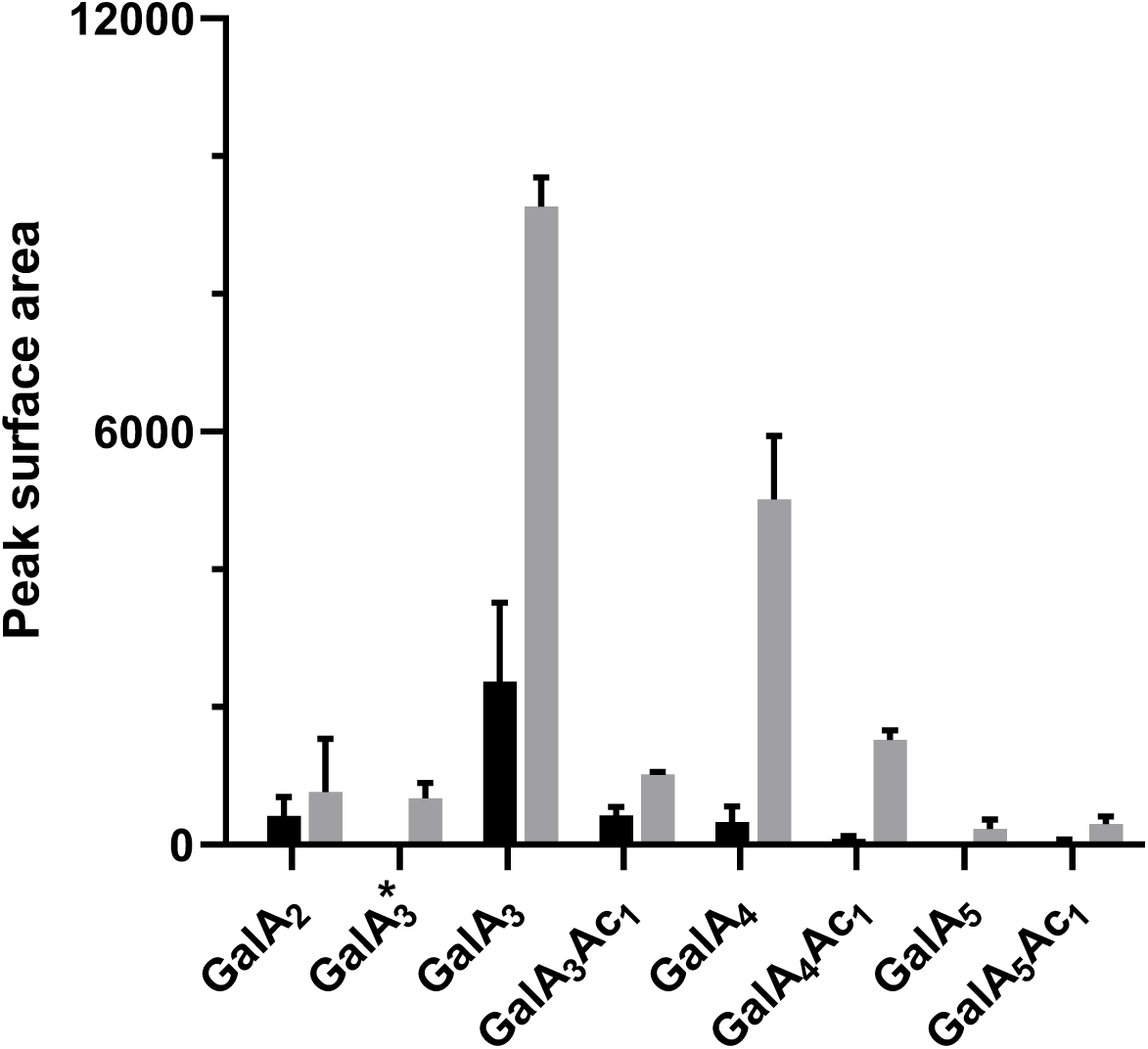
Analysis of OGs released by VdPelB from flax roots. VdPelB was incubated overnight with roots from Évéa (spring flax, partially resistant to Verticillium wilt, black) and Violin (winter flax, more susceptible to Verticillium wilt, grey). Values correspond to means ± SE of three replicates. Subscript numbers indicate the DP, DM and DA.

## 4. Conclusion

We have fully characterized, by multidisciplinary approaches, a novel pectinolytic enzyme from *V.dahliae*, VdPelB, that belongs to the PLL family. The protein was crystallised and its 3D structure determined at a high resolution. VdPelB showed a conserved structure, with typical topology for PLs and the active site harboured three conserved Asp coordinating Ca^2+^ and Arg involved in the β-elimination mechanism. The binding groove of VdPelB showed conserved aa that are characteristic of PLs, with MD simulations confirming the lower dynamics/higher affinity of the enzyme towards non-methylesterifed pectins. As inferred from structural and dynamical analyses, VdPelB showed high activity on non-methylesterified substrates, with a maximum activity at pH 8 and 35°C. The analysis of the structure led the identification, in the VdPelB, of peculiar aa that are normally present in PNL. In particular, G125R mutant showed increased activity on PGA and switch in pH optimum from 8 to 9. The analysis of the digestion products of showed that VdPelB act as an endo enzyme and that it can release a large diversity of OGs with a preference for non-methylesterified and acetylated products. The OGs generated by VdPelB from Verticillium-partially tolerant and Verticillium-sensitive flax cultivars showed that the enzyme could be a determinant of pathogenicity, as a function of pectins’ structure.

## Acknowledgements

We wish to thank Sylvain Lecomte and Mehdi Cherkaoui for providing the *Verticillium dahliae* DNA, Martin Savko and the staff at Proxima 2a beamline (Synchrotron SOLEIL, Gif sur Yvette, France) for X-ray diffraction and data collection.

## Author’s contribution

**Josip Safran**: Conceptualization, Data curation, Formal analysis, Investigation, Methodology, Writing - original draft. **Vanessa Ung**: Data curation, Investigation, Methodology. **Julie Bouckaert**: Data curation, Investigation, Methodology. **Olivier Habrylo**: Formal analysis, Investigation, Methodology. **Roland Molinié**: Data curation, Formal analysis, Investigation, Methodology. **Jean-Xavier Fontaine**: Data curation, Formal analysis, Investigation, Methodology. **Adrien Lemaire**: Investigation, Methodology. **Aline Voxeur**: Investigation, Methodology. **Serge Pilard**: Investigation, Methodology. **Corinne Pau-Roblot** Conceptualization, Methodology. **Davide Mercadante**: Data curation, Methodology, Writing - review & editing. **Jérôme Pelloux**: Funding acquisition, Conceptualization, Writing - review & editing. **Fabien Sénéchal**: Conceptualization, Writing - review & editing.

## Supplementary materials caption

**Table S1.**
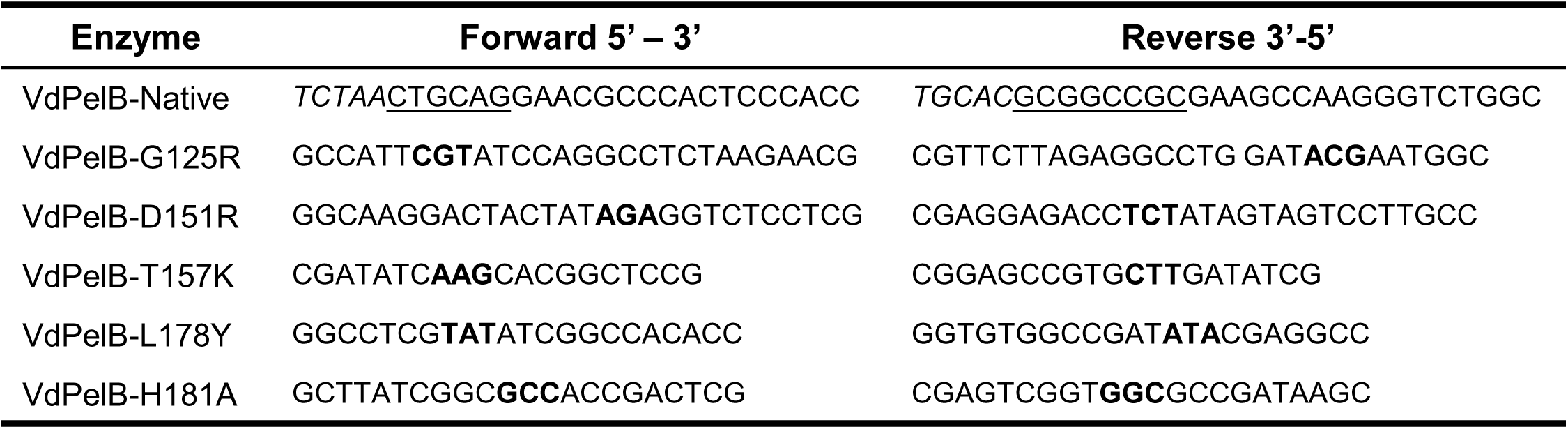
Primers for cloning mutated forms of VdPelB into pPICzαB expression vectors. Restriction enzymes sites for PstI and NotI are <u>underlined</u> added bases are written in italics. Mutation bases are **bolded.**

**Table S2.**
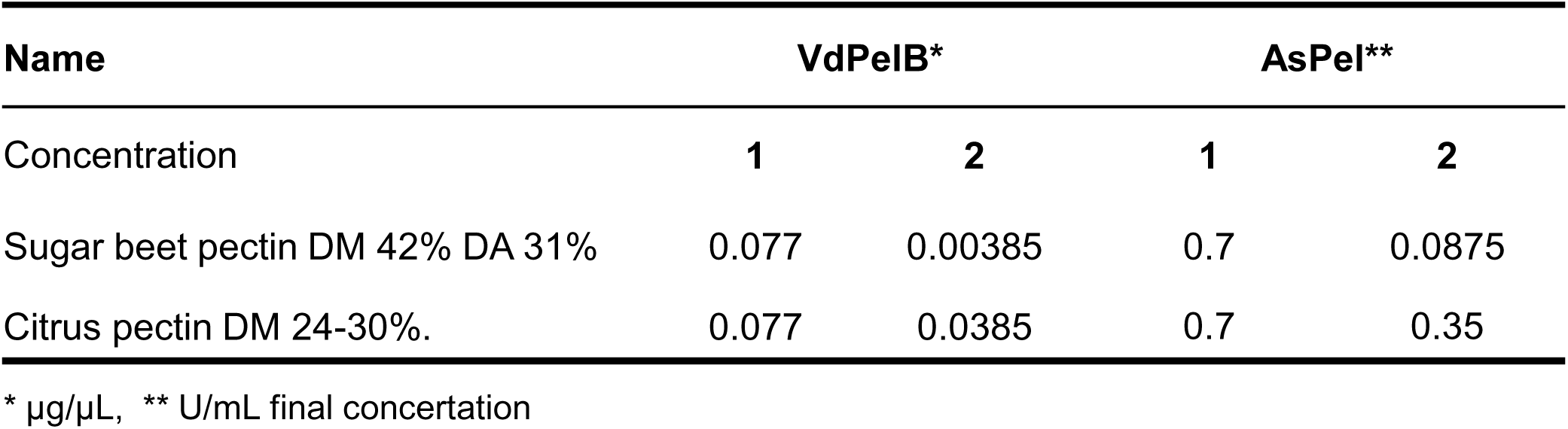
VdPelB and AsPel concentration used for pectin degradation and OGs analysis. The different enzymes concentrations were used to have enzymes at iso-activities.

**Fig. S1.**
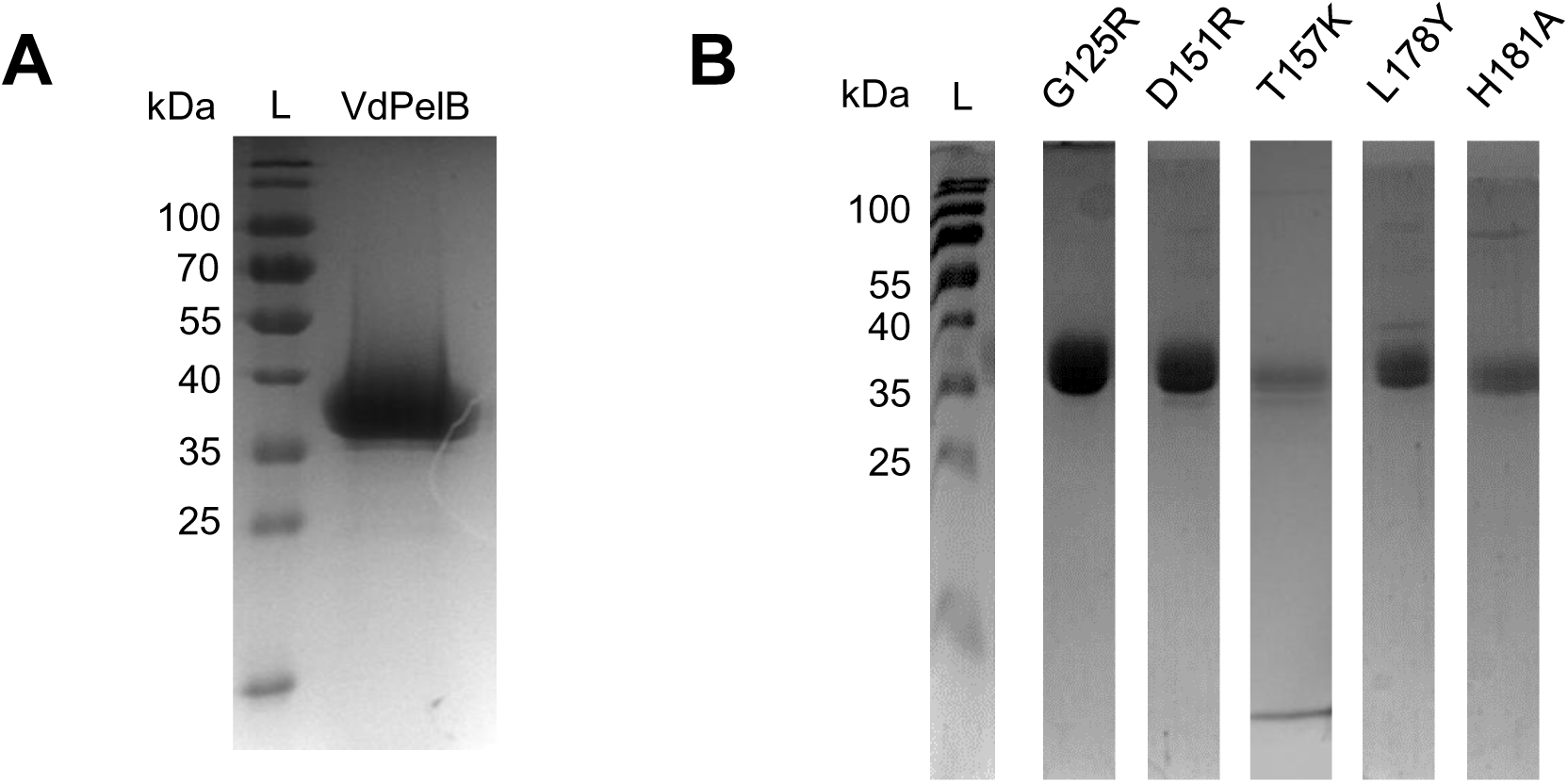
Purification and SDS-PAGE analysis of wild-type and mutated forms of VdPelB. Wild-type (A) and mutated forms (B) of VdPelB were purified using 1 mL Ni NTA colon. Proteins were resolved on a 12% polyacrylamide gel (SDS-PAGE) and were stained by Coomassie blue. L-ladder. The figure of SDS-PAGE representing mutants (B) was composed of several different images.

**Fig. S2.**
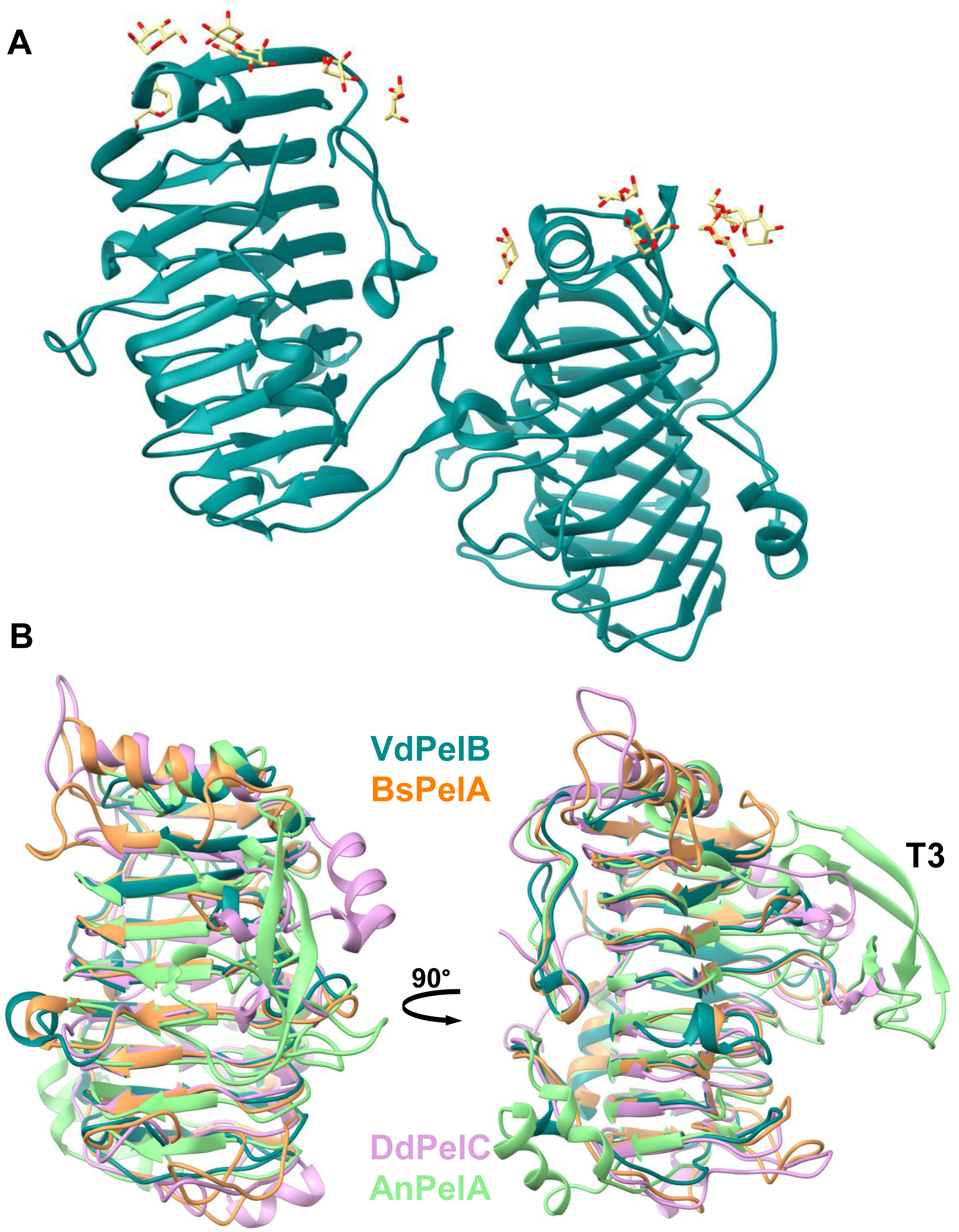
VdPelB crystal packing and structural alignment. A) Ribbon diagram of two molecules of VdPelB crystalized in P1 21 1 space group. **B)** Structural alignement of VdPelB with BsPelA, DdPelC, AnPelA. VdPelB shares highest structural alignment with BsPelA (PDB: 3VMV, orange) with 30.06% sequence identity and rmsd of 1.202 Å. Second best alignment was with DdPelC (PDB: 1AIR, plum) with 24.20% identity and rmsd of 1.453 Å. Pectin lyase withT3 loop described in AnPelA (PDB: 1IDJ, green).

**Fig. S3.**
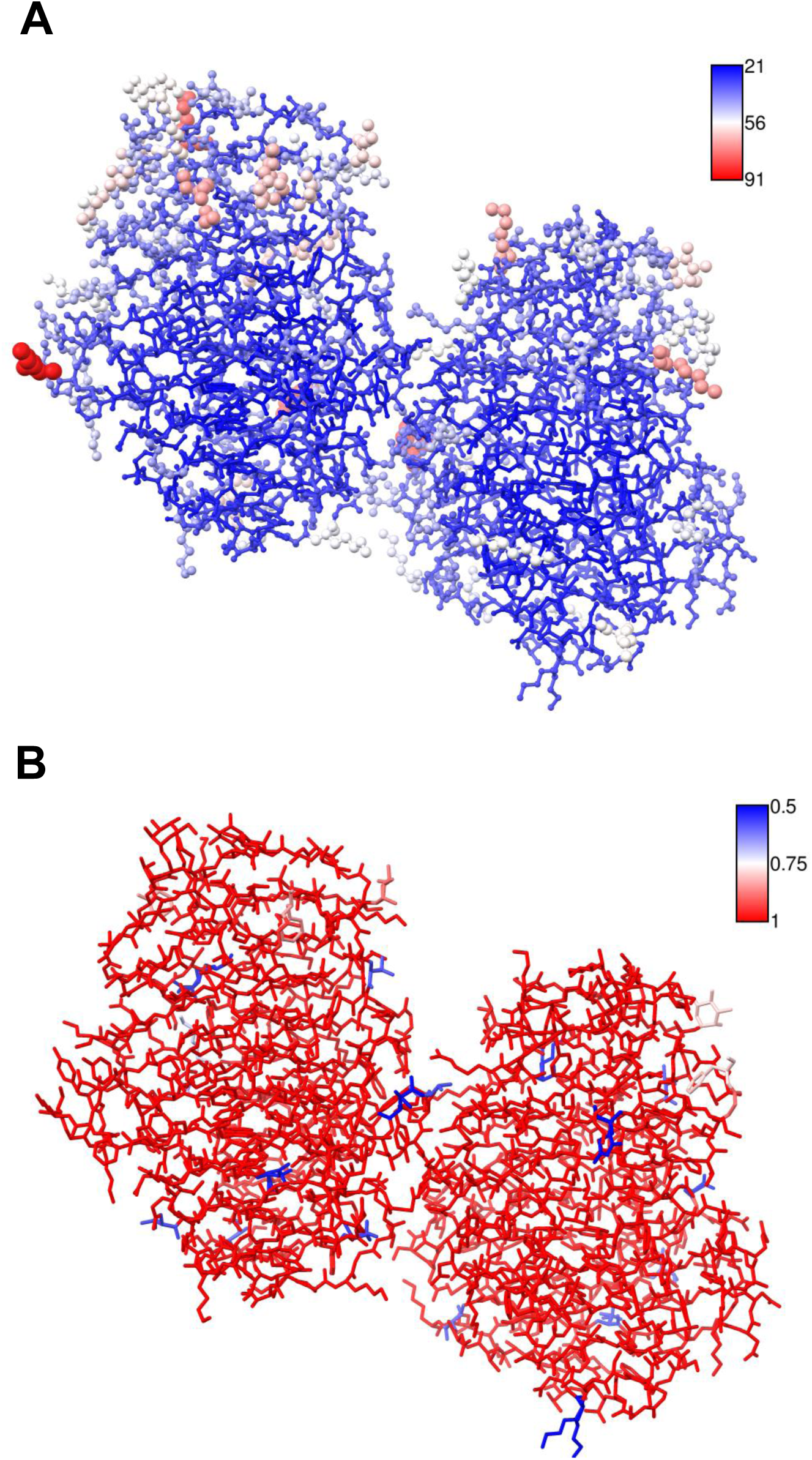
Average B-factors and occupancies of VdPelB. A) VdPelB chain A and B colored by B-factors. **B)** VdPelB chain A and B colored by occupancies.

**Fig. S4.**
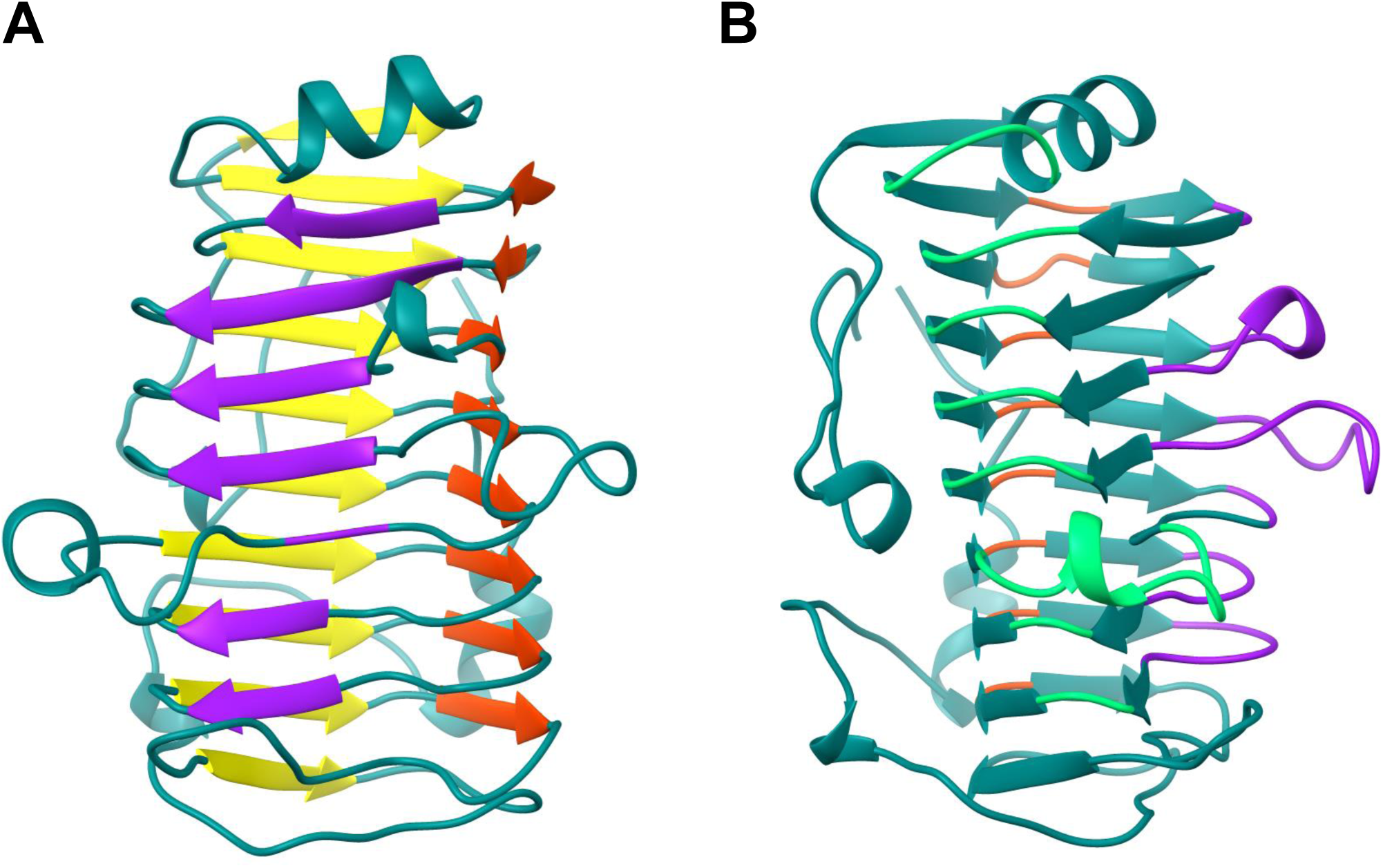
β-sheets and T-turns structures of VdPelB. A) Ribbon structure representing β-sheets (PB1-purple, PB2-yellow and PB3-red) B) Ribbon structure representing T-turns for VdPelB (T1-lime green, T2-orange red, T3 medium purple. β-strands and T-turns are named accordingly to Petersen et al. 1997.

**Fig. S5.**
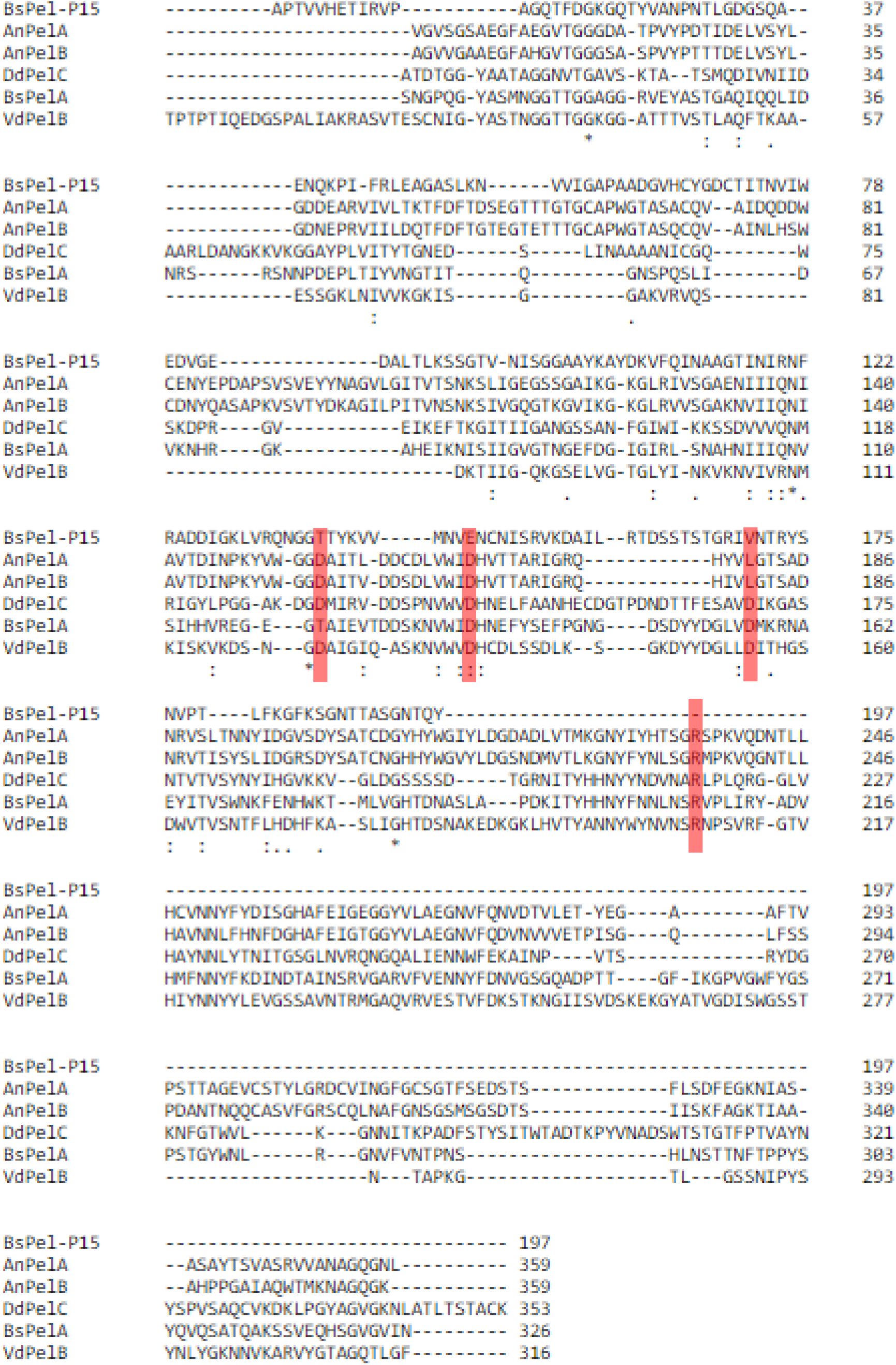
Primary sequence alignement of VdPelB with BsPelA, BsPelA-P15, DdPelC, AnPelA, AnPelB. VdPelB primary sequence alignement with *Dickeya dadanti* DdPelC (P11073), Bacillus Sp. KSM-P15 BsPel-P15 (Q9RHW0), Bacillus sp. N16-5 BsPelA (D0VP31), *Aspergillus niger* AnPelA (Q01172), AnPelB (Q00205). Conserved AA are red boxed.

**Fig. S6.**
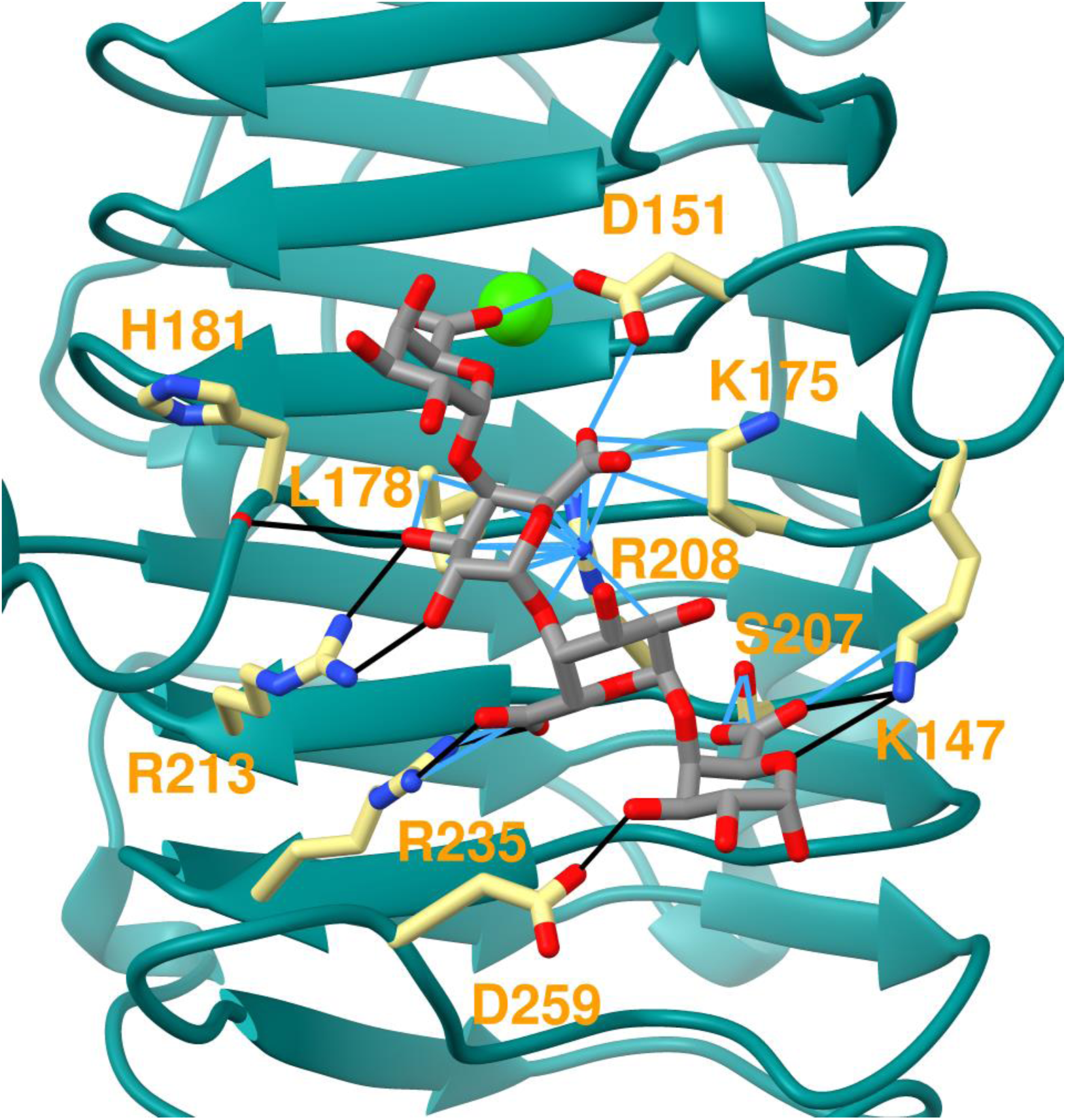
Superimposed tetramer ligand from DdPelC to VdPelB structure. Tetramer ligand from DdPelC (PDB: 2EWE, gray) superimposed to VdPelB structure. Aa involved in interaction are yellow-coloured. Hydrogen bonds and Van der Waals contacts are coloured black and blue.

**Fig. S7.**
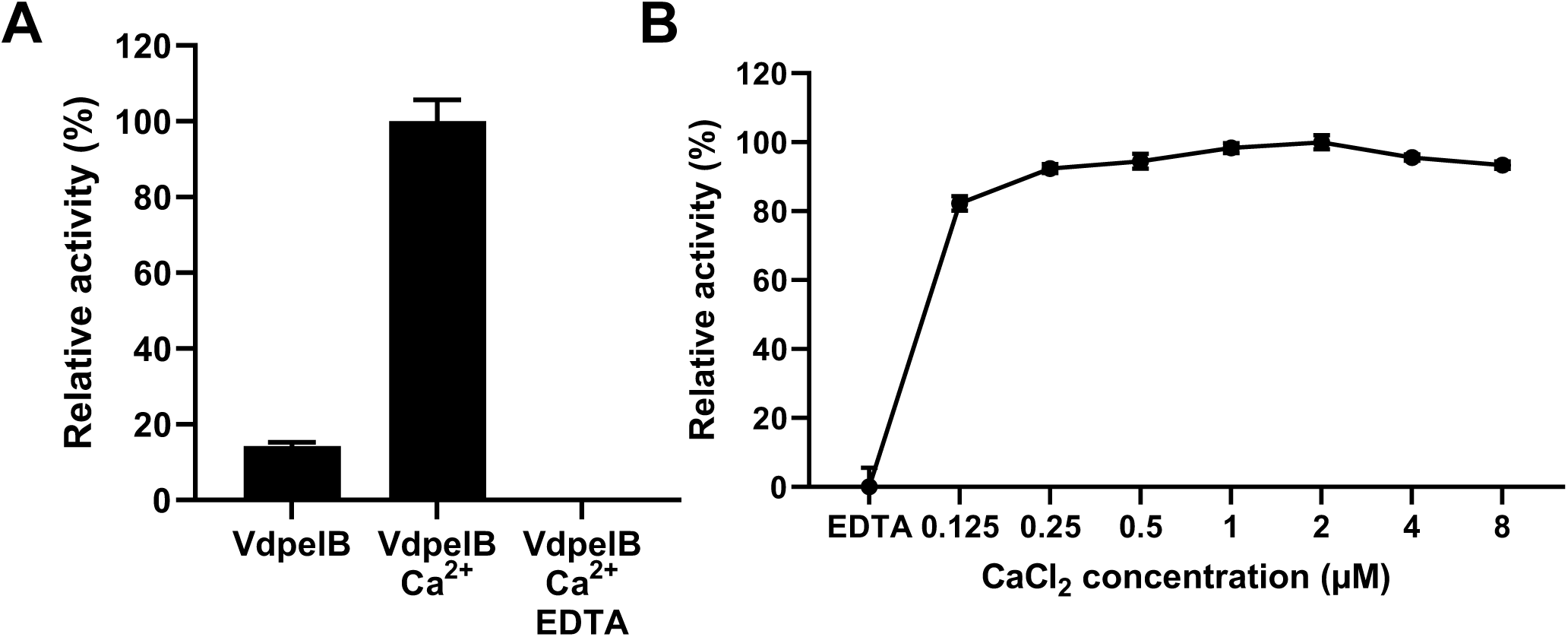
Effects of Ca^2+^ on VdPelB activity. A) Calcium dependence of VdPelB activity. The activities were measured after 12 min of incubation with PGA, with/without addition of Ca^2+^or Ca^2+^ and EDTA, at 35°C B) PGA digested using VdPelB for 12 min at 40°C and pH 8 with increasing concentrations of Ca^2+^.

**Fig. S8.**
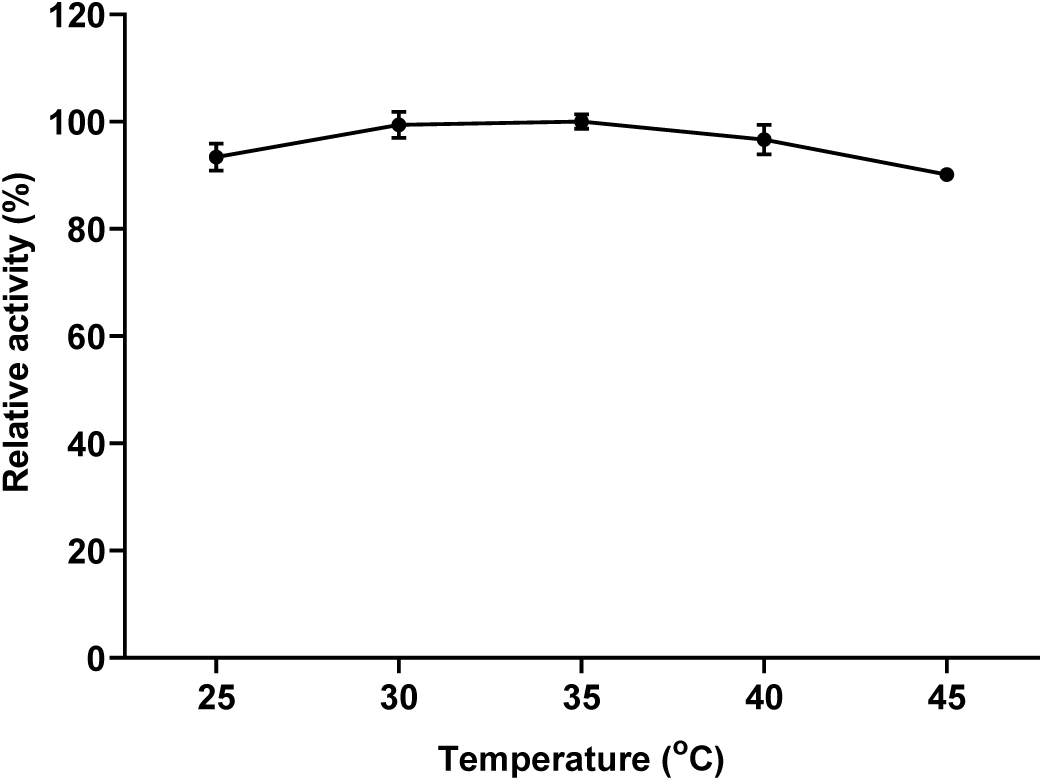
Temperature-dependence of VdPelB-N activity. The activities were measured after 8 min of incubation with PGA at pH 8. Values correspond to means of three replicates.

**Fig. S9.**
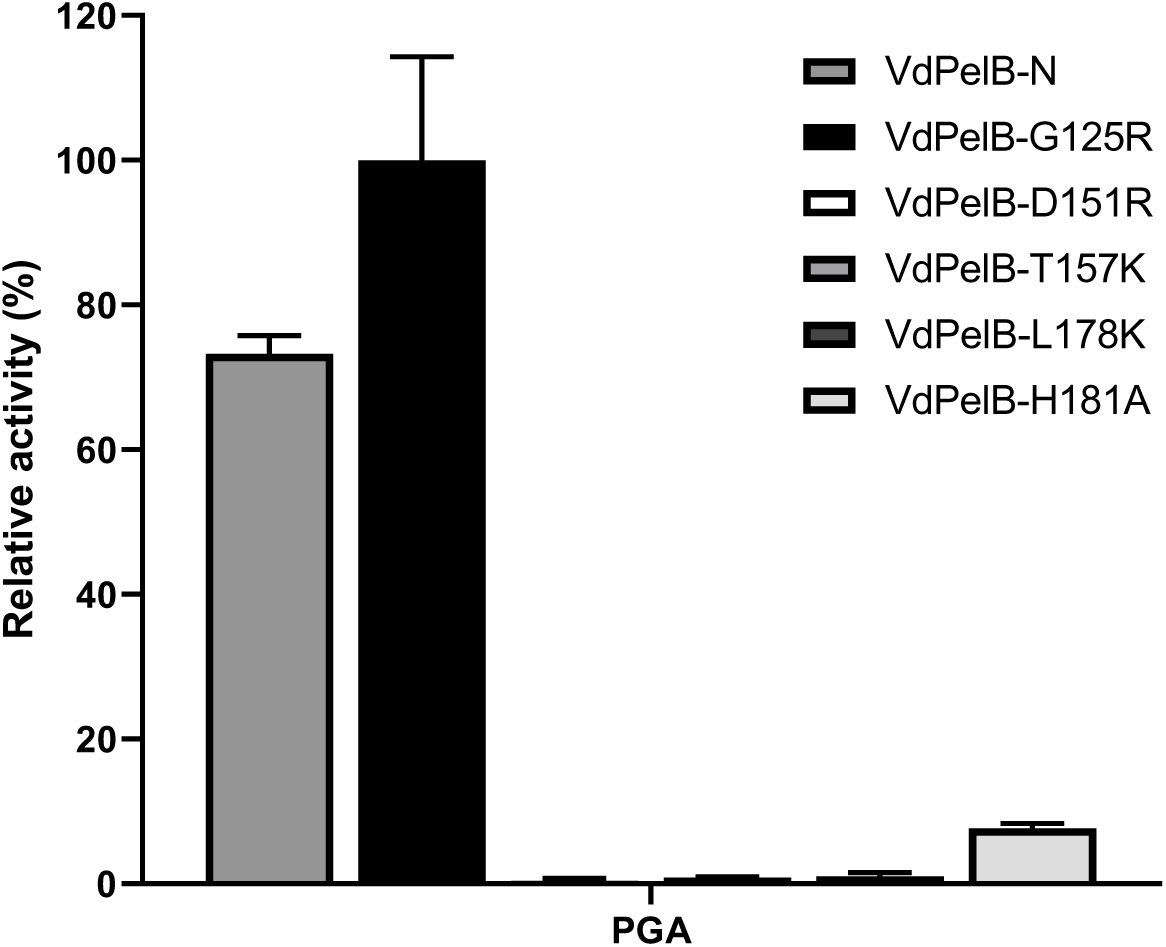
VdPelB mutants activity on PGA. The activities were measured after 12 min of incubation with PGA with addition of Ca^2+^ at 35°C. Values correspond to means of three replicates ± SD.

**Fig. S10.**
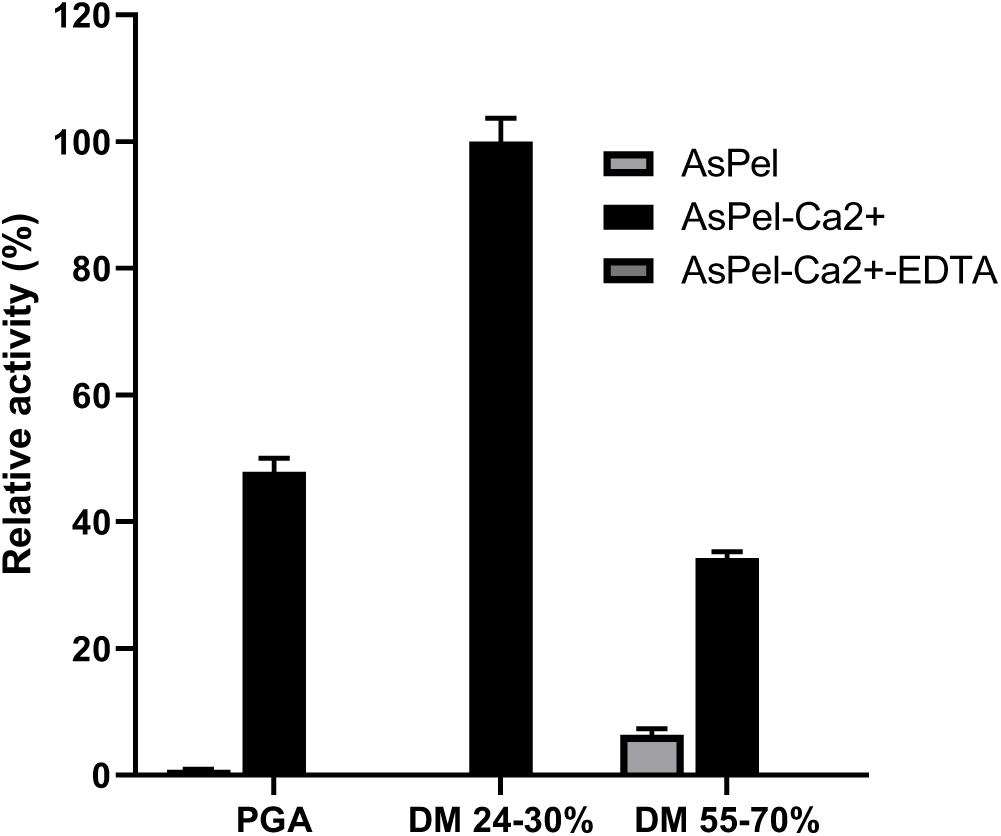
Substrate-dependence of AsPel. The activities were measured after 12 min of incubation with PGA, pectins DM 24-30%, DM 55-70%, with/without addition of Ca^2+^or Ca^2+^ and EDTA, at 35°C. Values correspond to means of three replicates ± SD.

**Fig. S11:**
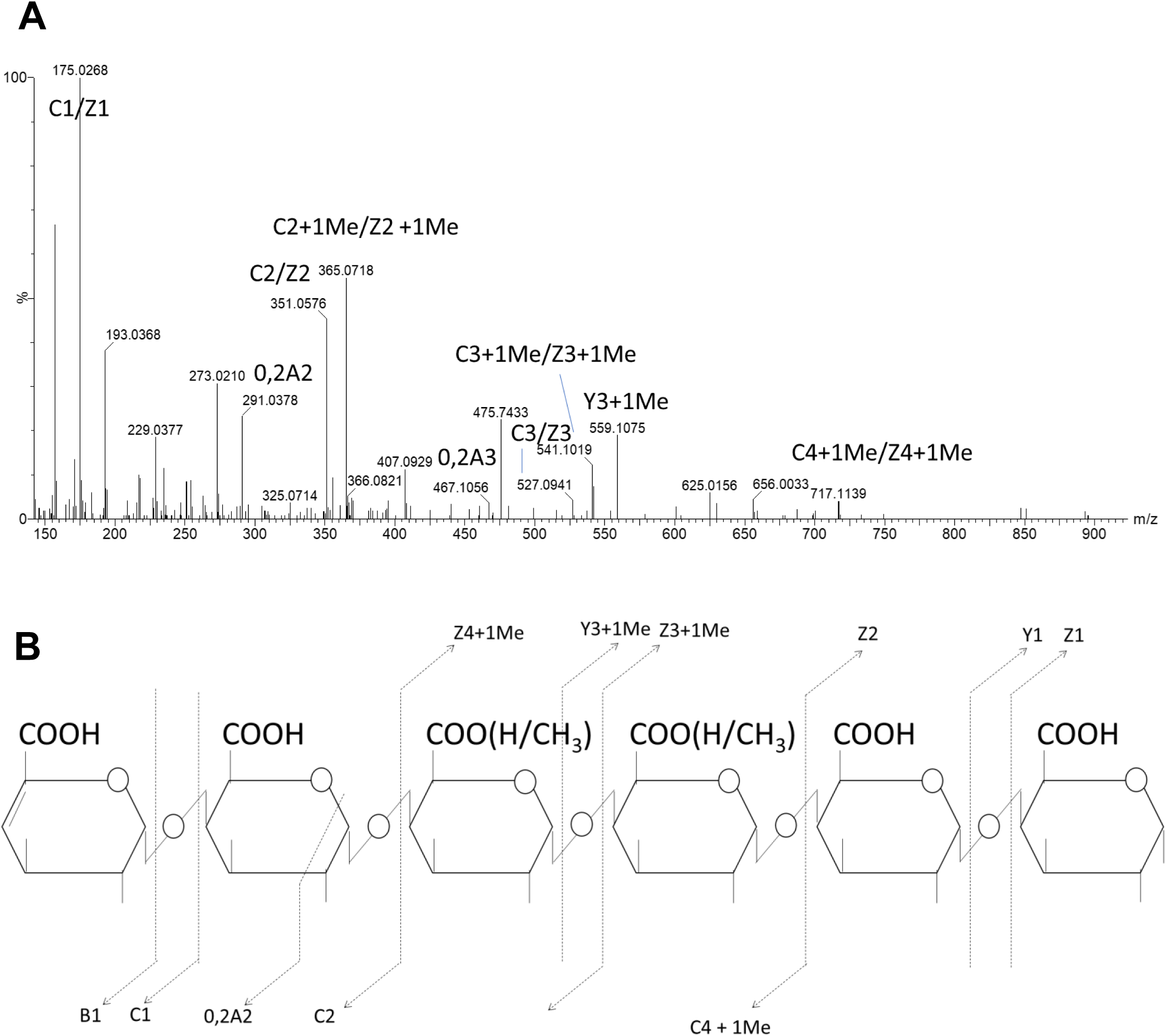
Example of MS^2^ fragmentation pattern of GalA_6_Me_1_-H_2_0. MS^2^ fragmentation pattern of GalA_6_-H_2_0 oligomer (m/z 1055,185252, *527,0889878*) produced by VdPelB from pectin DM >85%. Subscript numbers indicate the degree of polymerization.

**Fig. S12:**
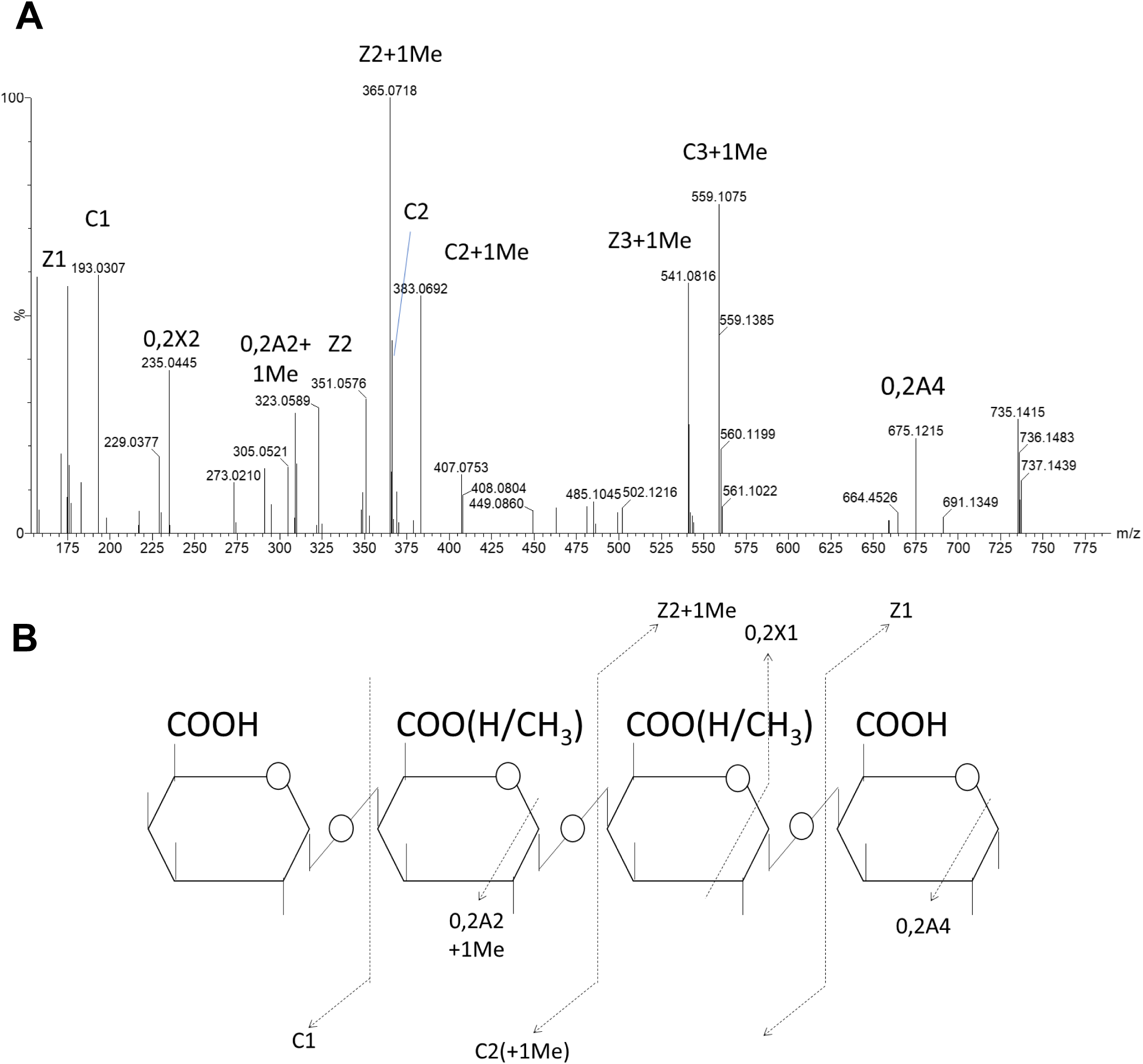
Example of MS^2^ fragmentation pattern of GalA_4_Me_1_. MS^2^ fragmentation pattern of GalA_4_Me**_1_** oligomer (m/z 735,1472907) produced by VdPelB from pectin DM >85%. Subscript numbers indicate the degree of polymerization.

